# Organization and triggered release of liposomes with DNA-based synthetic condensates

**DOI:** 10.1101/2025.09.27.678970

**Authors:** Diana A. Tanase, Layla Malouf, Roger Rubio-Sánchez, Catherine Fan, Karan Jain, Bortolo Matteo Mognetti, Lorenzo Di Michele

## Abstract

Cells use a combination of membrane-bound and membrane-less compartments to dynamically orchestrate internal biochemical processes and sustain intracellular communication. Recapitulating the hierarchical integration and interplay between these physically and chemically diverse structures is required to enhance the functionalities of synthetic cells and other advanced biomimetic systems. Here, we describe the use of synthetic DNA condensates to selectively uptake and spatially organize lipid vesicles, interacting with the condensates thanks to cholesterol-DNA anchors. By modulating anchor density, the liposomes can be programmably localized on the surface or interior of the condensates, while base-pairing selectivity can be leveraged to target individual internal domains in multi-phasic condensates. The embedded liposomes can be released by adding a nucleic acid trigger and captured by a second condensate population, thus imitating extracellular vesicles in their ability to support long-range cellular communication. This modular platform demonstrates the potential of DNA-based condensates to program the spatial distribution of membranous subcompartments and to support dynamic cargo-handling capabilities. These features are valuable for engineering cell mimics, microreactors, and delivery systems.

---

Cells orchestrate their complex biochemical processes using a diverse array of dynamic, self-organizing compartments. Alongside well-known membrane-bound organelles, ^1^ membrane-less organelles (MLOs) formed via liquid–liquid phase separation (LLPS) of proteins and nucleic acids are increasingly recognized as key regulators of cellular function. ^2–6^ MLOs recruit biomolecules to control processes such as stress signaling, gene regulation, and immune sensing, ^2–6^ making them attractive therapeutic targets.^7,8^ The structure and function of both membrane-bound and membraneless compartments are highly dynamic.^9^ LLPS enables the reversible formation and dissolution of MLOs to facilitate rapid adaptation and homeostasis,^10,11^ while membranous organelles are constantly reshaped, for instance to generate, traffic, and degrade vesicles. These include intracellular compartments such as endosomes, lysosomes and Golgi-derived vesicles,^1,12^ and extracellular vesicles that mediate long-range communication.^13,14^ Besides simply coexisting, membrane-bound and membraneless compartments can be hierarchically nested, with condensates embedding smaller membranous structures within their interior. ^9^ The clustering of synaptic vesicles, for instance, is thought to be mediated by their embedding within Synapsin 1 condensates, ^15,16^ while Balbiani bodies are membrane-less organelles that pack membrane-bound organelles including mitochondria, the ER, and Golgi. ^17^

A central goal of bottom-up synthetic biology is to recreate hierarchical and compartmentalized architectures to engineer both living and synthetic cells. ^18–22^

Remarkable progress towards this goal has been made for both membrane-enclosed and membrane-less compartmentalization. Giant Unilamellar Vesicles (GUVs) serve as plasma membrane mimics, with bulk and microfluidic methods developed to tune their properties and build multi-vesicle architectures.^23–27^ Encapsulation of smaller liposomes within larger vesicles has also enabled the construction of nested multiscale assemblies.^28–30^ In parallel, synthetic MLOs have been generated from proteins,^31–34^ RNA,^35–38^ or DNA.^39,40^ Nucleic acids, in particular, enable the construction of multi-domain condensates leveraging selective base-pairing,^36,37,41–44^ steric effects, ^45^ or reaction networks.^46–48^ These architectures support spatially organized internal functionality such as computation, ^49–51^ transcription,^45,46,52^ and enzymatic pathways. ^47,53,54^ Synthetic nucleic acid condensates have been deployed as model systems to study the biophysics of LLPS, leveraging the precise control that nucleic acids afford over valency and interaction strength, which are harder to program with alternative condensate-forming building blocks.

Hybrid architectures combining membranebound and membrane-less compartments have typically relied on GUVs encapsulating condensates or hydrogel networks.^55–59^ Current methods, however, do not enable the precise control over the relative spatial organization of nested and co-existing membranous and membraneless compartments we observe in biological cells. Similarly, challenges remain with inducing dynamic rearrangements of the microcompartmentalized architectures. Both these features would be highly valuable for engineering advanced biochemical pathways in synthetic cell mimics.

Here we introduce a hybrid platform that enables membranous compartments to be programmably captured and localized within DNA-based condensate scaffolds, imitating natural vesicle-in-condensate architectures.^9,15–17^ By tuning DNA–membrane affinity, the location of liposomes can be continuously shifted from the surface of condensates to their interiors, as we demonstrate experimentally and rationalize with theoretical modeling. Base-pairing selectivity allows vesicles to be targeted to specific condensate populations or phases,^43^ while toehold-mediated strand displacement^60–62^ enables their triggered release, imitating the production of extracellular vesicles by living cells.

We demonstrate that these *synthetic extracellular vesicles* (sEVs), released by *sender* condensates, can be captured by distinct *receiver* condensates, ^45,63,64^ imitating cellular pathways for long-range material exchange and communication. Finally, we show that vesicle-decorated condensates can be uptaken by live cells, laying the foundations for future intracellular applications. This modular DNA-liposome framework thus offers a powerful route for programming the spatial distribution of membranous compartments within synthetic condensates and their subsequent triggered release, opening opportunities in the design of synthetic cells, organelles, and delivery platforms.

## Results and Discussion

### Design of Hybrid DNA-Liposome Condensates

In Fig. 1, we summarize the design principles underpinning hybrid DNA–liposome condensates, comprising Large Unilamellar lipid Vesicles (LUVs) and condensate-forming DNA nanostructures. The latter include DNA junctions with four 20 base pair (bp) doublestranded (ds)DNA arms, dubbed “nanostars” (NSs), and 20 bp duplex “linkers”.^43^ All NS arms and both ends of the linkers feature 6 nucleotide (nt) single-stranded (ss) DNA sticky ends (SEs), with SEs on the NSs being complementary to those on linkers. As such, linkers act as mediators of NS-NS interactions, leading to condensate formation. ^43^

**Figure 1:**
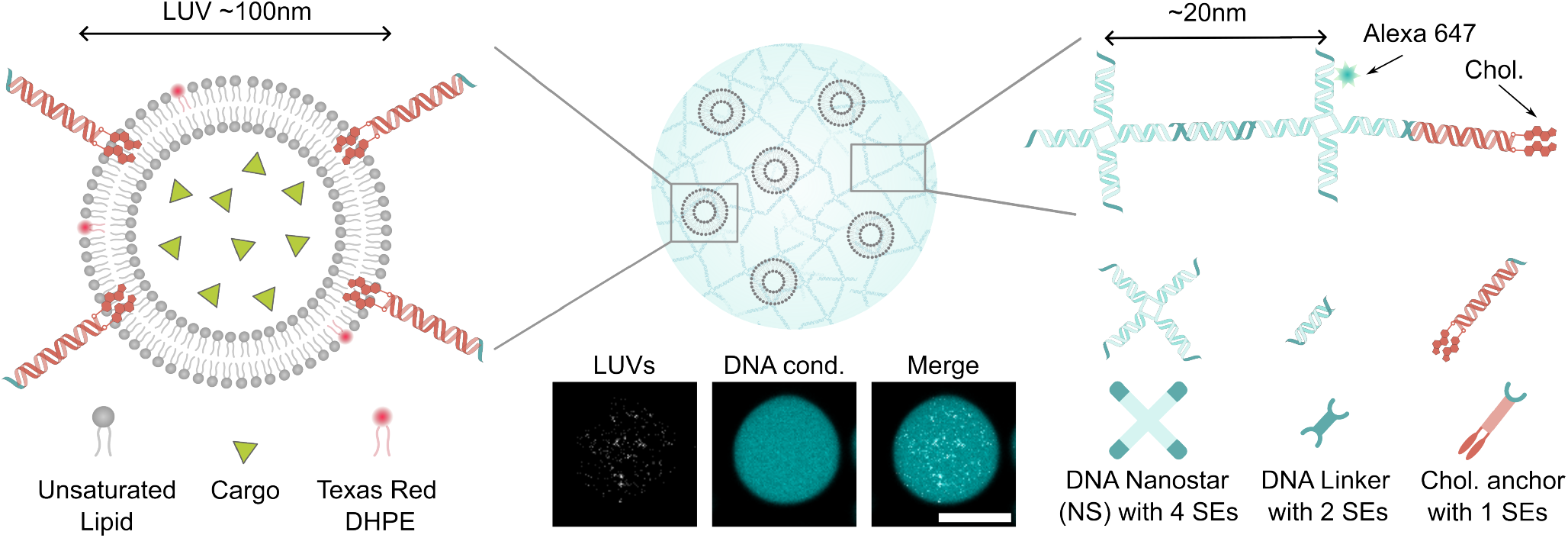
Assembly principles for hybrid DNA–liposome condensates. Schematic of Large Unilamellar lipid Vesicles (LUVs) embedded within DNA condensates via cholesterol-mediated anchoring (not to scale). LUVs, which may include encapsulated cargo, are pre-incubated with double-cholesterol DNA anchors and co-annealed with condensate-forming DNA nanostars (NS) and duplex DNA linkers. NSs and linkers interact selectively through complementary sticky ends (SE), leading to condensation. A single-stranded overhang on the cholesterol-modified DNA anchor is complementary to target the NS sticky-end, enabling the incorporation of liposomes within the DNA network. Confocal micrographs (bottom) show liposomes labeled with Texas Red DHPE (gray), DNA condensates labeled with Alexa 647 (cyan) by replacing a fraction of the NS forming strands with its fluorescently labeled version, and the composite image. Scale bar: 10 µm.

The size of the NSs and linkers, and the length of the SEs, are chosen to ensure that the building blocks remain stable at temperatures above the condensate melting point. Specifically, the 20-bp double-stranded domains in the NSs and linkers disassemble at *T*_m_ > 65^*◦*^C, well above the melting temperature of the condensates (*T*_m,cond_ ≈ 50^*◦*^C).^43^ This design facilitates a robust two-step assembly protocol: stocks of NSs and linkers are pre-annealed and mixed with LUVs at room temperature, followed by a brief heating step to 55^*◦*^C, sufficient to melt the SE bonds and ensure uniform mixing, but low enough to preserve the structural integrity of the building blocks. Additionally, the 6-nt SEs (Table S1) ensure that the condensates remain stable at or slightly above physiological temperatures. Finally, we note that while longer NS arms and linkers would be compatible with these constraints, building larger constructs would require longer individual DNA strands, which are more prone to truncation defects arising during solid-phase synthesis.

The linker:NS concentration ratio is fixed to 2:1, to ensure matching stoichiometry between SEs on the two species. LUVs, with nominal diameter of ∼ 100 nm (Fig. S1) are prepared via extrusion using 99 mol % 1,2-dioleoylsn-glycero-3-phosphocholine (DOPC), doped with 1 mol % Texas Red 1,2-dihexadecanoyl-sn-glycero-3-phosphoethanolamine, triethylammonium salt (Texas Red DHPE). To control liposome-condensate interactions, LUVs are functionalized with 36 bp dsDNA “anchors”, labeled with two cholesterol moieties at one end and a ssDNA SE (6 nt) at the other end.^65–68^ Cholesterol moieties ensure stable membrane insertion of the anchors,^69,70^ while the SE, complementary to those on NSs, mediates affinity for the DNA condensates. See Tables S1 and S2 for DNA sequences and Methods for details on liposome and nanostructure preparation.

One-pot co-annealing of DNA NSs (0.5 µM), linkers (1 µM), and functionalized liposomes, causes the SEs on NSs to hybridize with complementary SEs on both the linkers and the liposome-bound DNA anchors, yielding hybrid DNA-liposome condensates (Fig. 1). Confocal microscopy (Fig. 1 and large fields of view in Fig. S2) confirms successful hybrid assembly, showing embedding of the LUVs (Texas Red DHPE) within the DNA condensates (Alexa 647). Figure S3 confirms that the fluorescent dye calcein, encapsulated in the liposomes, is retained during the co-annealing process and can be subsequently released by disrupting the LUVs with Triton X-100 surfactant.

### Programming the Spatial Distribution of Liposomes in Hybrid Condensates

Having demonstrated the construction of hybrid DNA condensates with embedded LUVs, in Figure 2 we explore the effect of changing the number of cholesterol-DNA anchors on the spatial distribution of liposomes. Throughout this work, we focus primarily on two conditions: LUVs with a higher anchor density (∼ 1250 anchors per vesicle, or ≃ 3.1 mol% cholesterol:lipids) and lower anchor density (∼ 125 anchors per vesicle or ≃ 0.3 mol% cholesterol:lipids) (Fig. 2**a**). In the high anchor coverage scenario, we aim to work at cholesterol anchor densities close to membrane saturation.^71^ See Supplementary Note I for details on estimating anchor coverage. Dynamic light scattering confirms a slight difference in hydrodynamic diameter of the LUVs between the two conditions, consistent with changing anchor coverage (Fig. S1).

**Figure 2:**
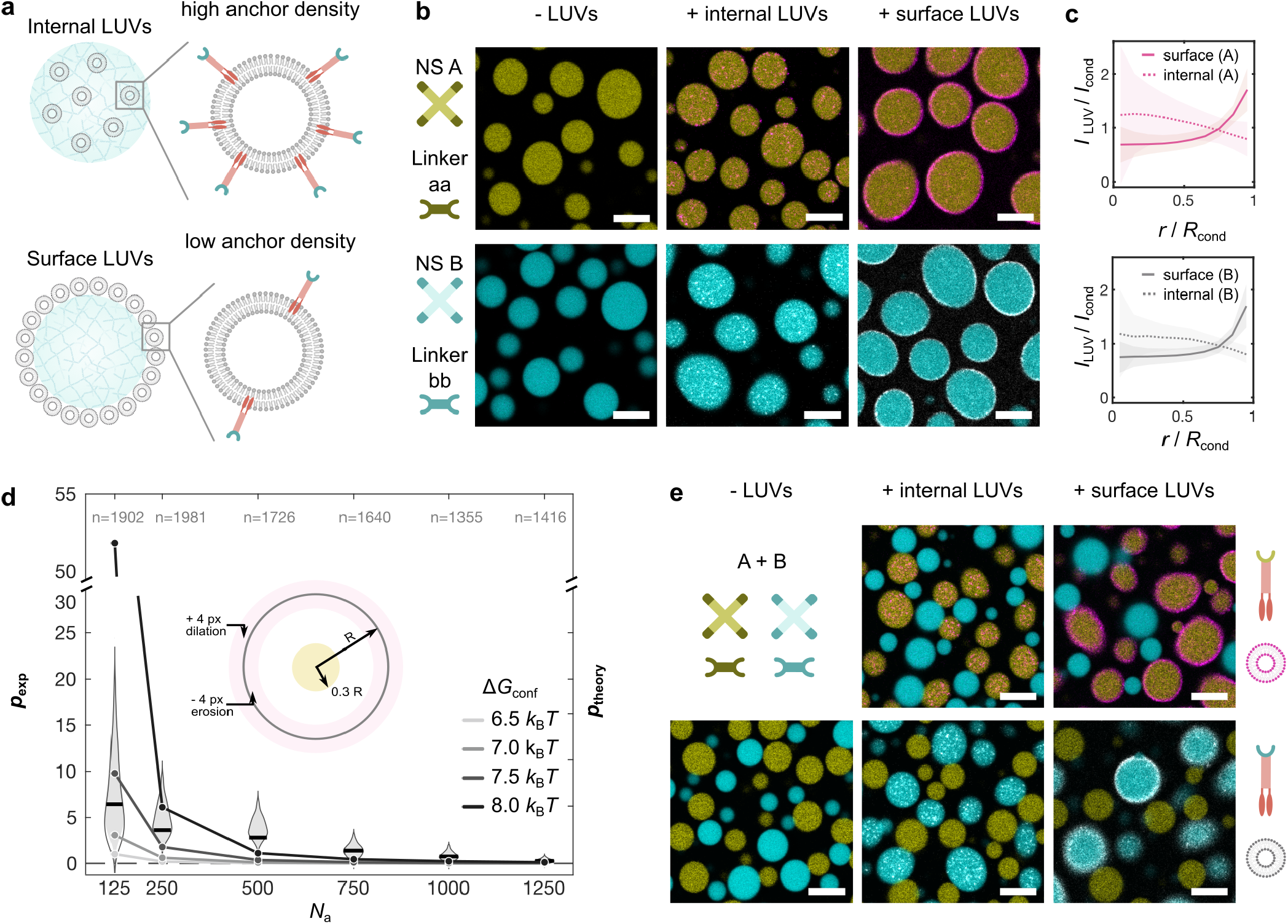
Programmable and targeted spatial distribution of LUVs in orthogonal DNA condensates. **a** LUVs can be programmed for internal sequestration within DNA condensates when decorated with a high density of DNA-cholesterol anchors (∼ 10^3^ anchors per vesicle, top), or for surface association when fewer anchors are included (∼ 10^2^ anchors per vesicle, bottom). Schematic illustrations are not to scale. **b** Confocal micrographs of orthogonal condensates A (top, yellow, labeled with Atto 488) and B (bottom, cyan, labeled with Alexa 647), assembled from stoichiometric mixtures of A NS + aa linkers and B NSs + bb linkers, respectively. Micrographs show representative examples of condensates without LUVs (left), with internally sequestered LUVs (center), and with surface-targeting LUVs (right). Liposomes target the two condensate types thanks to overhangs complementary to either NS A (yellow) or NS B (cyan). All LUVs share the same composition but differ in the cholesterol anchors used, which selectively target either A or B condensates. LUVs targeting NS A are shown in magenta (Texas Red DHPE–labeled), and those targeting NS B are shown in gray (Texas Red DHPE–labeled). **c** Radial intensity profile of LUV fluorescence, *I*_LUV_, normalized by the DNA signal, *I*_cond_, relative to the condensate centroid. Data correspond to panel b (top: condensates made with NSs A; bottom: condensates made with NSs B). Solid lines show mean *I*_LUV_*/I*_cond_ across individual condensates ± standard deviation as a function of normalized radial distance, *r/R*_cond_, with *R*_cond_ the condensate radius. Over 1000 condensates of each type were analyzed across multiple fields of view. Segmentation and analysis are detailed in the Methods section, SI. **d** Transition from surface-bound to internalized LUVs as a function of cholesterol anchors per LUV, *N*_a_. Experimental data (left axis) show *p*_exp_, namely the intensity ratio of surface to core fluorescence in condensates made with NSs A and linkers aa (see main text). The right axis shows theoretical predictions of the probability ratio between surface localization and embedding, *p*_theory_ (see main text). The two limiting cases of *N*_a_ = 125 anchors (surface-localized) and *N*_a_ = 1250 (fully engulfed) were used throughout this work. For experimental measurements, representative examples are shown in Fig. S4 and were prepared in triplicate. The number of analyzed condensates (*n*) is indicated above each corresponding violin plot, alongside the median value. **e** In mixed samples of co-existing A (yellow, Atto 488) and B (cyan, Alexa 647) condensates, LUVs exhibit selective targeting based on the sequence of the cholesterol anchors SEs. All scale bars: 10 µm.

As summarized in Fig. 2**b**, we proceed to integrate high- and low-anchor density LUVs in DNA condensates. We use two sequenceorthogonal DNA condensates, one constructed with “A” NSs (Atto 488, yellow) and associated “aa” linkers and the second consisting of “B” NSs (Alexa 647, cyan) and “bb” linkers. ^43^ Using anchors that selectively interact with the relevant A or B NSs we observe that, for both condensate types, anchor density controls the spatial distribution of liposomes: LUVs with a high anchor density are embedded in the bulk of the condensates, while vesicles with lower density of DNA anchors accumulate at the surface of condensates. Analysis of radial intensity profiles in Fig. 2**c** confirms the visual assessment of LUV distribution in Fig. 2**b**, showing near uniform distribution across the condensates for high-anchor-density LUVs and a substantial accumulation at the surface of condensates for LUVs with sparser DNA coating. This degree of LUV localization mirrors the organization of membranous compartments at the surface or within the interior of biological condensates. ^9^ The observation that LUV localization is dependent on anchor density is consistent with recent literature having reported that the ability of condensates to uptake particles depends on the affinity between particles and the condensed phase, with weakly-binding particles partitioning on the condensate surface and strongly-binding ones being uptaken. ^72^

In our system, tuning anchor density or, in other words, the multivalent avidity of the LUVs, offers a simple route to controlling LUVcondensate interactions without having to alter the affinity of individual anchor-condensate interactions, which would require redesigning the SEs or anchors. As shown in Fig. 2**d** and Fig. S4, we leverage the precise control afforded by this strategy to explore how the LUV distribution changes with anchor density for values between the highand low-coverage conditions tested above. We quantify partitioning of the LUVs between the surface and core of the condensates from confocal micrographs. We sample LUV fluorescence, *I*_LUV_, within a thin shell at the condensate surface and within its core. To correct for optical projection artifacts, we normalize *I*_LUV_ by the signal sampled in the DNA fluorescence channel, *I*_DNA_, measured within the same regions (Methods section, SI). Due to the finite lateral resolution of the confocal microscope, we are unlikely to be able to distinguish LUVs adsorbed at the condensate surface from those embedded just beneath it. Therefore, we expect (*I*_LUV_*/I*_DNA_)_surface_ − (*I*_LUV_*/I*_DNA_)_core_ to be roughly proportional to the excess (volume) density of surface-adsorbed versus embedded LUVs. We can then define a normalized partitioning parameter *p*_exp_ = (*I*_LUV_*/I*_DNA_)_surface_*/*(*I*_LUV_*/I*_DNA_)_core_ − 1, which quantifies the excess accumulation of LUVs at the condensate surface relative to its core.

Large *p*_exp_ values are observed at low number of anchors per LUV, *N*_a_, indicating surface accumulation. *p*_exp_ then progressively decreases as *N*_a_ increases, converging towards 0 for *N*_a_ ≳ 1000, indicative of no excess surface accumulation or, in other words, a uniform distribution of LUVs throughout the condensates. For any given *N*_a_, *p*_exp_ shows a broad distribution across the condensate population, which we ascribe largely to statistical noise due to the relatively small number of LUVs sampled within the segmented areas. We further note that *p*_exp_ changes smoothly with *N*_a_, in contrast to previous reports showing a sharp transition between surface binding and condensate engulfment for particles with variable binding affinity for the condensates. ^72^

To rationalize the experimental observations on the effect of anchor coverage on LUV distribution, we constructed a simple model, detailed in Supplementary Note II. Our calculations account for the multivalent nature of LUV-condensate interactions and the lateral mobility of the anchors enabled by membrane fluidity, both of which may influence the adhesion and uptake processes. Briefly, LUVs are modeled as hard spheres of radius *R*, decorated with membrane-bound anchors. Building on an established framework, ^73^ anchors are considered as rigid rods of length *L* = 12.2 nm (36 bp), which can freely pivot and diffuse laterally. The exposed condensate surface is assumed to contain unpaired SEs at a fixed surface density, 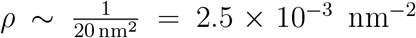, estimated from the condensate mesh size. Anchors and unpaired NS arms can bind through their complementary SEs, assumed point-like and fully flexible, with interaction free energy Δ*G*_0_ + Δ*G*_conf_, where Δ*G*_0_ is the SE hybridization free energy, estimated as ≃ − 12 *k*_B_*T* using NUPACK^74^ for A-type SE sequences, and Δ*G*_conf_ is a term accounting for configurational effects.^73^ Under these assumptions, we can compute a multivalent LUV-condensate interaction free energy, *F* (*h*), where *h* is the distance between the condensate surface and the center of the vesicle. This can be compounded with the surface-energy contribution associated with deformation of the DNA phase as *f* (*h*) = *F* (*h*) + 2*πγR*(*R* − *h*), where *γ* ∼ 10^−6^ N m^−1^ ^75^ is the condensate surface tension. Minimizing *f* (*h*) allows us to determine the equilibrium penetration depth of the LUV, *h*_eq_.

As shown in Fig. S5, for a fixed *R, h*_eq_ displays a sharp transition from surface-adsorbed to embedded states with increasing number of anchors, which contrasts with the smooth trend observed experimentally (Fig. 2**d**). However, we note that the experimental LUV population is polydisperse in size (Fig. S1) and likely also in anchor surface density. To account for these sources of polydispersity, for each nominal value of *N*_a_, we compute *h*_eq_ for a polydisperse sample of 4000 LUVs with radii drawn from a log-normal distribution fitted to DLS data (Fig. S1 and Supplementary Note II) and numbers of anchors drawn from a Poisson distribution with average equal to *N*_a_. We then compute the ratio *p*_theory_ between the number of surface-bound and embedded LUVs in the polydisperse sample, which can be compared with the experimental surface-partitioning parameter, *p*_exp_ (Fig. 2**d**).

As a result of sample polydispersity, *p*_theory_ qualitatively reproduces the smooth trend observed in *p*_exp_, increasing with decreasing *N*_a_ and converging towards 0 for *N*_a_ ≳ 1000. Agreement between *p*_theory_ and *p*_exp_ is maximized for values of Δ*G*_conf_ between 6.5 *k*_B_*T* and 8 *k*_B_*T* . This range is reasonable given the interpretation of Δ*G*_conf_ as an effective penalty term accounting for constraints beyond the simple hybridization free energy, such as configurational restrictions at the nanostar network level, reduced accessibility of binding sites, or steric effects arising from junction stiffness.

We note that, while our model captures the experimental trends, full quantitative agreement is not expected due to effects that cannot be captured by our simple framework. These include, for instance, LUV deformability and non-spherical shape, which are likely to vary substantially across the population due to variability in excess area and the presence of multilamellar constructs. ^76,77^

While our equilibrium model is able to qualitatively predict LUV embedding within the condensates for sufficiently high anchor coverage, we point out that the co-annealing protocol we use to prepare the hybrid condensates is likely essential for enabling internalization. The internal dynamics of NS phases slow down with decreasing temperature following an Arrhenius relationship, ^78^ ultimately resulting in gelation at sufficiently low temperatures.^78–80^ During the annealing process, condensates nucleate, grow and readily coalesce, as we have previously characterized in the absence of LUVs.^43^ In this phase, LUVs would be able to access the interior of the condensate while it remains in a low-viscosity state. As the temperature decreases, the exponential increase in viscosity leads to the immobilization of LUVs within the DNA network. Consistent with this hypothesis, earlier studies on interactions between LUVs and cholesterolized DNA nanostar networks showed that LUVs added post-assembly decorate the condensate surface rather than penetrate its interior, despite LUV-condensate interactions likely being stronger than those in the present study.^45^

Finally, we tested LUV targeting selectivity by co-annealing samples of A-type and B-type condensates in equimolar amounts, with LUVs decorated with anchors complementary to only one of the relevant linkers (Fig. 2**e**). We observe that LUV uptake or surface decoration occurs only for the target condensate types, consistent with the expected bond selectivity.

### Spatial Patterning of Liposomes in Condensates with Tunable Phase Mixing

Having demonstrated control over the spatial distribution of LUVs in individual condensates, alongside targeted complexation in two-condensate mixtures, in Figure 3 we proceed to engineer liposome arrangement in more sophisticated multi-phase architectures.

**Figure 3:**
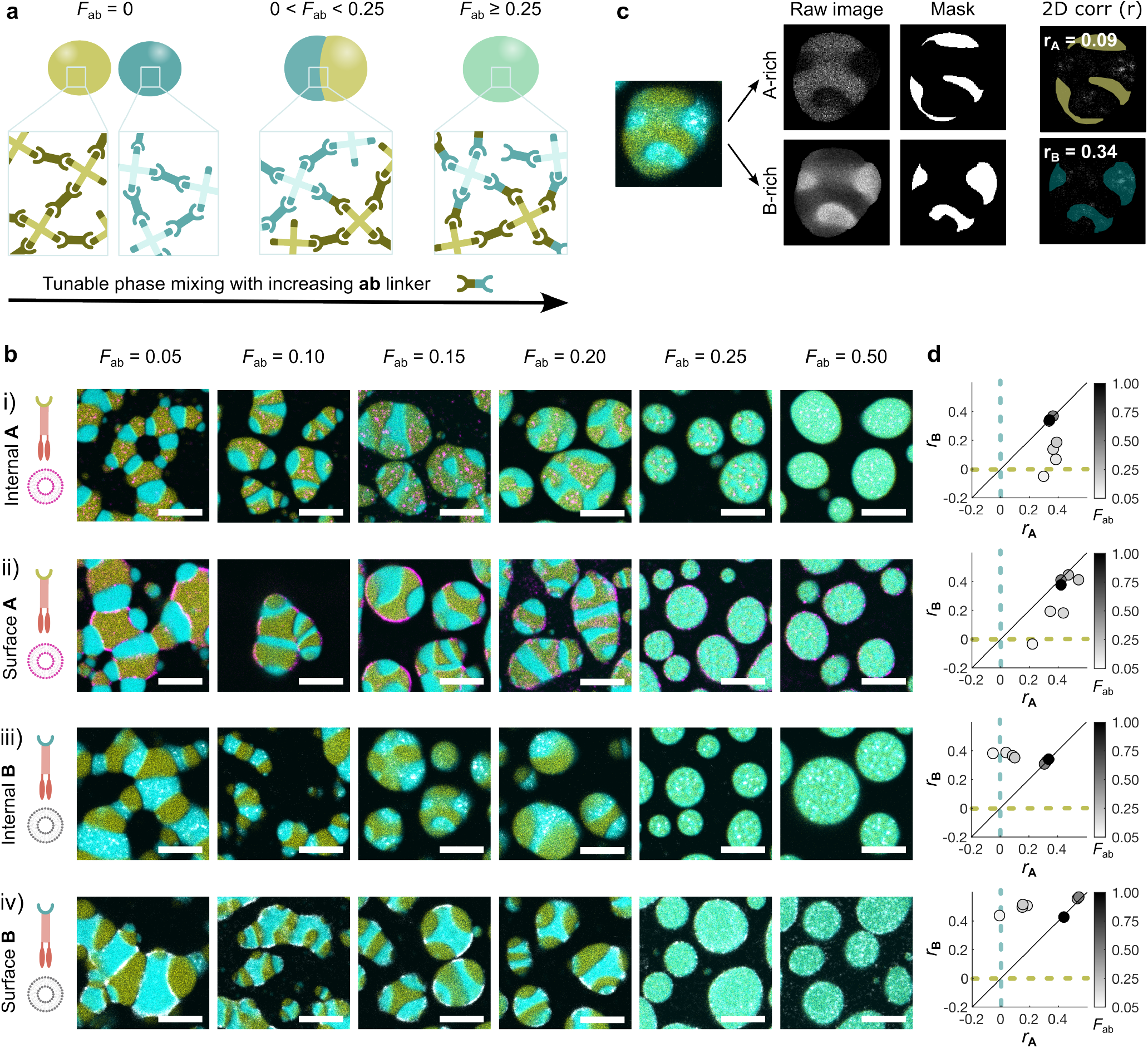
Liposome patterning in binary DNA condensates with tunable phase mixing. **a** Including a cross-binding linker, ab, which connects NSs A and B, enables the formation of biphasic condensates with tunable interphase mixing. ^43^ As the fraction of the ab linker, *F*_ab_ = [ab]*/*([aa] + [ab] + [bb]) increases, interphase mixing increases, with fully mixed single-phase condensates emerging at sufficiently high *F*_ab_. **b** Confocal micrographs (contrast adjusted from the lowest to the highest pixel intensity values) of condensates with increasing *F*_ab_, with either the A-rich (yellow, Atto 488) or B-rich (cyan, Alexa 647) phases targeted by internally-sequestered or surface associated LUVs (magenta or gray, Texas Red DHPE). **i)** Internally-sequestered LUVs in the A-rich phase. **ii)** Surface-tethered LUVs in the A-rich phase. **iii)** Internally-sequestered LUVs in the B-rich phase. **iv)** Surfacetethered LUVs in the B-rich phase. Grayscale micrographs of individual channels are shown in Figs S6-S9 for better visualization. **c** Confocal micrographs and segmentation steps for a hybrid DNA-liposome condensate (*F*_ab_ = 0.15) with LUVs internally-sequestered in the B-rich phase. To quantify the co-localization of LUVs with either phase, the raw images in each channel are segmented to identify corresponding masks. Then, the 2D correlation coefficient is calculated between the fluorescence signal of LUVs and either the A-phase mask (*r*_A_) or the B-phase mask (*r*_B_), condensate by condensate. **d** Scatter plot of the median values of 2D correlation coefficients (*r*_B_ vs *r*_A_) calculated over the condensate populations for samples shown in panel c. Stronger LUV localization in the target phases is associated with larger *r*_A_ or *r*_B_ values. *F*_ab_ is specified by the color bar on the right. The number of condensates analyzed and associated median absolute deviation values are provided in Table S3. All scale bars: 10 µm.

In a recent contribution, we showed that introducing a third linker, “ab”, bridging NSs A and B, enables control over the formation of biphasic condensates and allows tuning of the interfacial tension between the A-rich and B-rich phases.^43^ For low fractions of the ab linker (*F*_ab_ = [ab]*/*([aa] + [bb] + [ab])), the condensates display coexistence of A-rich and B-rich internal domains, while fully mixed, monophasic condensates emerge at higher *F*_ab_ (Fig. 3**a**). Here, we use equal and fixed concentrations of A and B NSs and, as for the case of pure condensates, stoichiometric proportions between complementary SEs are imposed, namely 4[A]=2[aa]+[ab] and 4[B]=2[bb]+[ab], with [aa]=[bb]. Hence, when increasing [ab], [aa] and [bb] are decreased accordingly.

As summarized in Fig. 3**b**, we then combined biphasic architectures with high- and low-DNA coverage LUVs, targeting individual A- or B- rich domains, demonstrating additional control over LUV organization. At low *F*_ab_, LUVs exhibit strong spatial colocalization with their target phase, and display the designed surface anchoring or engulfment behaviors, as observed in single-phase condensates (Fig. 2). As *F*_ab_ increases, the distinction between the two phases diminishes, resulting in more uniform LUV distributions across the condensate. Grayscale images are provided in Figs S6-S9 for the individual channels merged to obtain the composite micrographs in Fig. 3**b** to more clearly distinguish the signals of the DNA phases and LUVs.

Quantitative image analysis supports these observations, with Fig. 3**c** summarizing the segmentation pipeline for condensate analysis. We segment the confocal images and calculate 2D correlation coefficients between the fluorescent signal of LUVs and that of both the Aand Brich DNA phases, *r*_A_ and *r*_B_ respectively. Scatter plots of median values of *r*_B_ against *r*_A_ show that higher phase segregation (low *F*_ab_) yields higher correlations between LUVs and their target phase, and lower correlation with the orthogonal phase (Fig. 3**d**, Table S3). As *F*_ab_ increases (darker shades of gray in Fig. 3**d**), median correlation values converge, indicating reduced phase specificity of LUV association. As expected for fully mixed condensates (*F*_ab_ ≥ 0.25), the median correlation values lie on the diagonal of the scatter plots.

To assess whether LUVs influence domain morphologies and partitioning, we performed a control experiment in which samples were annealed without LUVs (Fig. S10). DNA condensates with and without LUVs display broadly similar morphologies. However, at *F*_ab_ = 0.20, the presence of LUVs leads to enhanced phase separation, as quantified in Fig. S11. In the absence of LUVs, condensates with these compositions exhibit irregular internal domains and an overall spherical shape, consistent with vanishingly small interfacial tensions between A-rich and B-rich regions.

De-mixed condensates both with and without LUVs display qualitatively analogous but non-identical domain morphologies, as also seen in natural condensates such as the nucleolus,^4,6^ emerging from the interplay between interfacial tension minimization and the intrinsically slow relaxation kinetics of these DNA networks.^43^

To further expand the range of accessible hybrid condensate-vesicle architectures, we explored the possibility of using LUVs to directly act as mediators of the interactions between NSs A and B, in the absence of ab linkers. LUVs were prepared with equal amounts of the two types of cholesterol anchors, complementary to SEs on NS A and NS B. As shown in Fig. S12, upon incubation of these bi-functional LUVs with mixtures of the two NSs and associated aa or bb linkers, we observed the formation of condensate networks similar to those made with *F*_ab_ = 0.05 (Fig. 3**c**). Here, vesicles accumulate at the interfaces between Aand B-rich domains, thus demonstrating a spatial pattern distinct from the ones accessible with ab linkers and mono-functional LUVs targeting individual phases.

### Programmable Release and Capture of Liposomes

The ability of cells to reorganize both membranous and membrane-less compartments underpins many of their most intriguing responses. We therefore proceed to demonstrate stimulus-induced reorganization of the multi-domain liposome-DNA architectures. To this end, as introduced in Figure 4, we modify aa and bb linkers by extending one of the two constituent DNA strands with a 6 nt toehold domain. ^60–62^ Trigger strands, *α* for aa and *β* for bb, can thus break apart the respective target linkers, causing the disassembly of either the A-rich phase, the B-rich phase, or both phases, depending on which trigger, or combination of triggers, is added (Fig. 4). This mechanism for phase-targeted disassembly is validated using co-existing, single phase A and B condensates (*F*_ab_ = 0) with epifluorescence micrographs at key time points (Fig. S13). The functionality of the strand displacement reactions is additionally confirmed by agarose gel electrophoresis (Figs S14–15). Sequences for toehold-modified linkers are provided in Table S1.

**Figure 4:**
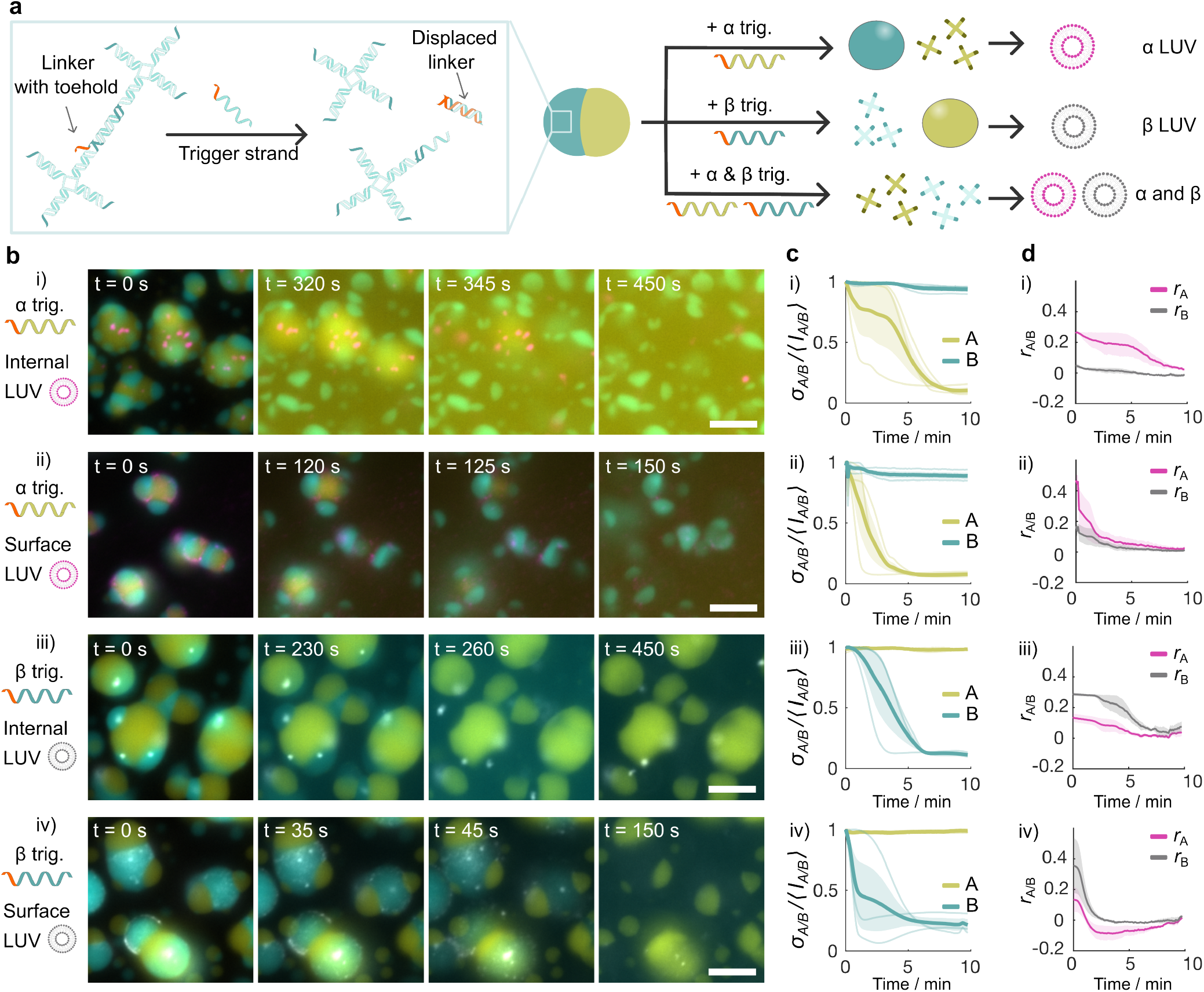
Sequence-specific release of LUVs from orthogonal DNA compartments. **a** DNA compartments in biphasic condensates are engineered to disassemble and release LUVs in response to sequence-specific trigger strands (*α* trig. and *β* trig.), via toehold-mediated strand displacement. Modified aa and bb linkers include a single-stranded toehold domain. Triggers are fully complementary to the toehold-bearing strand in the linkers. The strand displacement reaction disrupts the linkers producing two monovalent constructs unable to cross-link NSs, thus causing disassembly of the targeted condensate phase (A-rich or B-rich). The schematic highlights the mechanism for B NSs and bb linkers (cyan); the same strategy is applied to compartment A (yellow). **b** Epifluorescence micrographs at selected time points show the release of both internally sequestered and surface-tethered LUVs (Texas Red DHPE) from A-rich (yellow, Atto 488) and B-rich compartments (cyan, Alexa 647) in samples with *F*_ab_ = 0.05, upon addition of the relevant triggers. All images are scaled from minimum pixel intensity to 1.5 × the maximum value determined in the first frame post trigger strand addition for clearer visualization during the release. Note that micrographs are collected with epifluorescence microscopy to enable fast timelapse acquisition, resulting in reduced z-resolution compared to Figs 1-3. All scale bars: 10 µm. **c** Mean coefficient of variation (dark solid line) ± SEM (standard error of the mean, shaded area) of the fluorescence intensity, computed as the ratio between the standard deviation (*σ*_A*/*B_) and the mean of the pixel intensities (⟨*I*_A*/*B_⟩) in the fluorescence channels of A and B phases. Curves are normalized to their maximum values and measured across three independent repeats from raw (unscaled) micrographs. Light-colored traces show individual measurements. Each plot corresponds to the sample type shown in the adjacent row in panel b. **d** Mean correlation coefficients (*r*_A*/*B_, solid lines) ± SEM (shaded area) between LUV signal and the A-rich (magenta)/B-rich (gray) phase. Each plot corresponds to the sample type shown in the adjacent row in panel b.

Using the toehold-responsive linkers, we assemble biphasic condensates with LUVs targeting either the surface or the bulk of either the A-rich or B-rich phases, as shown in Fig. 4**b**. An ab linker fraction *F*_ab_ = 0.05 is chosen for these experiments, as this resulted in the most pronounced phase-selectivity for LUVs, whose signal cross-correlation was positive with the target phase and negative for the non-target phase (Fig. 3**d**). Figures S16–17 present larger fields of view for the samples in Fig. 4**b**, showing both the merged channels and the isolated LUV channel, which facilitates clearer visualization of the liposomes.

Upon addition of either the *α* or the *β* trigger strands, and consequent targeted phase disassembly, both internal and surface-tethered LUVs are selectively released from their respective target compartments, consistent with the expected orthogonality. Disassembly and release occurs within several minutes of trigger strand addition, although some variability in the onset of release is observed, likely due to local differences in trigger availability caused by manual mixing.

During condensate disassembly, we observe an increase in background fluorescence intensity in the channel corresponding to the phase being etched due to the local release of labeled strands before they diffuse away.

To quantify disassembly kinetics, we extract the time dependent normalized coefficient of variation in fluorescence intensity, *σ*_A*/*B_*/* ⟨*I*_A*/*B_⟩, for each of the two phases (A-rich or B-rich) and each sample type (Fig. 4**c**). *σ*_A*/*B_ and ⟨*I*_A*/*B_⟩ are the standard deviation and the mean of the pixel values, respectively, evaluated in the full field of view at each time point. *σ*_A*/*B_*/* ⟨*I*_A*/*B_⟩ reflects spatial heterogeneity in the fluorescence signal, which decreases as the targeted phase disassembles. As expected, the targeted compartments exhibit a marked decrease in *σ*_A*/*B_*/* ⟨*I*_A*/*B_⟩ over time, whereas the nontargeted compartments remain stable.

Visual inspection of the microscopy images confirms that LUVs initially localized to the domains targeted for disassembly are released in solution, imitating the generation of extracellular vesicles,^81^ including processes such as apoptosis, in which cells fragment into membranebound compartments that are released and captured by neighboring cells.^13,82^ LUV release is also reminiscent of synaptic vesicles being freed by the disassembly of synapsin condensates.^15,83^ Since individual LUVs are close to the diffraction-limited resolution of the microscope, and therefore difficult to track individually, we employed the same correlation analysis as in Fig. 3**c-d** to study the release process (Fig. 4**d**). Masks for the two DNA phases were defined in the first frame of the timelapse and applied to all subsequent frames (segmentation details are provided in the Methods section, SI). In all four sample types, the mean correlation between the LUV signal and the targeted phase (*r*_A*/*B_ for A-rich and B-rich, respectively) decreases as LUVs are released. For LUVs sequestered internally, the correlation with the non-targeted phase (Fig. 4**d**, B-rich in i or Arich in iii, respectively) remains roughly constant. By contrast, for surface-tethered LUVs (Fig. 4**d** ii and iv), the correlation with the nontargeted phase also decreases, although starting from a lower correlation value. We attribute this to LUVs that, while tethered to the targeted phase, are positioned at the interface between A-rich and B-rich domains. These orthogonal release experiments confirm that toehold-mediated strand displacement can selectively and rapidly (≤ 10 min) trigger the release of LUVs from specific DNA compartments.

Finally, we tested whether released LUVs could be recaptured by a separate class of DNA condensates, demonstrating a full release–recapture cycle within a synthetic system of membrane-less and membranous compartments. For this purpose, we use amphiphilic DNA nanostars in which SEs are replaced by terminal cholesterol modifications, previously shown to robustly condense upon thermal annealing due to cholesterol-cholesterol hydrophobic interactions (Fig. 5**a**).^63,64^ In these condensates, cholesterol moieties on the surface offer a natural mechanism for capturing the released LUVs (Fig. 5**a**).^45^ Note that, different from DNA nanostars interacting through SEs, amphiphilic nanostars can form solid, crystalline phases under suitable buffer conditions and if annealed sufficiently slowly, as previously reported and evident from the polyhedral shape of the condensates. ^63,64^ Oligonucleotide sequences used for the amphiphilic condensates are shown in Table S4.

**Figure 5:**
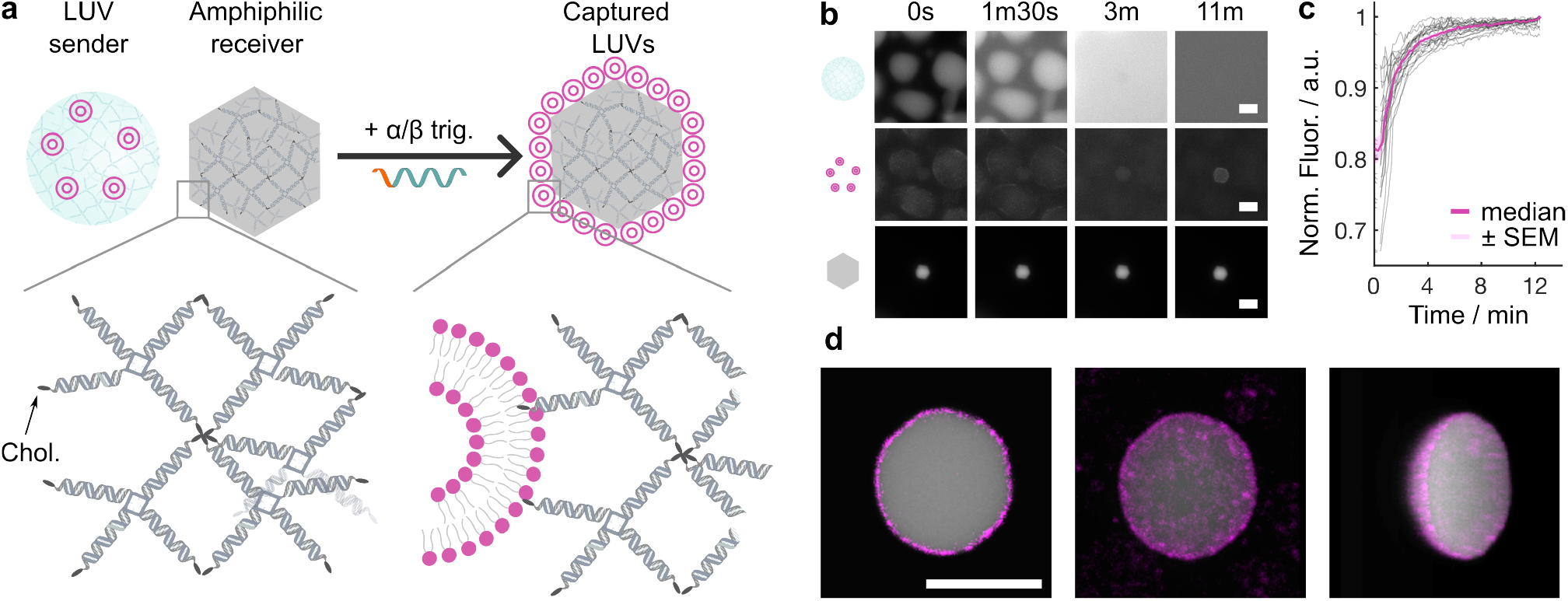
Release and capture of synthetic liposomes by amphiphilic condensates. **a** Schematic summarizing the release of LUVs from DNA condensates in response to a trigger strand (*α/β* trig.) and the subsequent capture of LUVs onto crystalline amphiphilic condensates via hydrophobic interactions (cholesterol molecules are depicted as dark gray ellipses). ^45,63,64^ **b** Epifluorescence micrographs (grayscale) at selected time points during LUV release and capture. Condensates (NS A, Atto 488) are shown in the top row, LUVs (Texas Red DHPE) in the middle row, and a representative amphiphilic condensate (Cy5) in the bottom row. Micrographs are scaled from minimum pixel intensity to 1.5 × the maximum pixel intensity as measured in the first frame after trigger strand addition. **c** Normalized LUV fluorescence intensity (Texas Red DHPE) relative to the maximum within each amphiphilic condensate ROI measured across full fields of view. Intensity increases over time as LUVs accumulate on the amphiphilic condensates. The magenta line shows the median, with the standard error of the mean shaded in light magenta. Grey lines show fluorescence signal traces for 21 ROIs. Additional independent measurements from different wells are shown in Fig. S19. **d** Confocal micrographs of an amphiphilic condensate (gray, Cy5-labeled) coated with captured LUVs (magenta, Texas Red DHPE-labeled). Shown are an equatorial slice (left), a 3D reconstruction—top view (center), and side view (right). LUVs were released from DNA condensates containing NS A (Atto 488) and surface-tethered LUVs (Texas Red DHPE). All scale bars: 10 µm.

We prepared TMSD-responsive A-type DNA condensates with surface-tethered LUVs, and added the relevant trigger strand in the presence of amphiphilic condensates (Cy5-labeled). Epifluorescence microscopy revealed progressive accumulation of LUV fluorescence on the amphiphilic condensates over time (Fig. 5**b**). Quantification of the normalized fluorescence intensity within ROIs corresponding to amphiphilic condensates (segmented as described in Methods section, SI and Fig. S18) confirms this accumulation, showing a clear increase over time following the addition of the trigger strand (Fig. 5**c**). In Fig. S19 we show analogous experimental results for three replicates of condensates made with either NS A or NS B, internally-sequestered or surface-tethered LUVs, demonstrating the robustness of the mechanism. LUVs released from condensates made with NS B (labeled with Alexa 647) were captured by fluorescein-labeled amphiphilic condensates (sequences in Table S5) to distinguish between the two fluorescent labels. We note that condensates with surface-tethered LUVs lead to a more significant release and subsequent capture by amphiphilic condensates, consistent with an easier release compared with internalized LUVs. An equivalent result is obtained in binary condensates (Fig. S20) made with *F*_ab_ = 0.05 and in which LUVs are released from the B-rich phase, although the monitoring of the release in epifluorescence microscopy is more challenging due to the increased number of fluorescent components.

To better visualize the LUVs captured by amphiphilic condensates, we acquired confocal micrographs, shown in Fig. 5**d** as equatorial slices and three-dimensional reconstructions. These images confirmed the successful accumulation of LUVs at the surfaces of amphiphilic condensates. Together, these results demonstrate that the released LUVs can be efficiently recaptured through hydrophobic interactions.

Our findings highlight the versatility of this programmable release-and-capture platform, which could find applications beyond the construction of synthetic cells, for instance, as payload delivery vehicles to living cells. To assess whether the hybrid condensates can be interfaced with biological cells, we performed a proof-of-concept experiment. Condensates (NSs A and linkers aa) were prepared with surface-localized LUVs (125 cholesterol-DNA anchors per vesicle) encapsulating calcein as a model cargo, and incubated with HEK293 cells (Fig. S21). Imaging immediately after addition and again 24 h later reveals that LUVs from the condensates are uptaken by the cells and remain detectable intracellularly at the 24 h timepoint. Incubation was carried out at 37 °C, a temperature at which condensates are expected to be more liquid-like, which may facilitate uptake. This observation demonstrates that the hybrid condensates can access an intracellular environment, establishing a first step towards biological applications. We emphasize, however, that developing these materials into fully functional delivery systems would require substantial further work, including a systematic characterization of the uptake mechanism and of intracellular cargo release, which lies beyond the scope of the present study. Such applications are nonetheless an increasingly active direction for synthetic condensates, which are being developed as platforms for intracellular delivery and sensing. ^84^ The ability of condensates to penetrate cell membranes, and the molecular determinants governing this process, are beginning to be elucidated for peptide- and RNA-based systems;^84^ our results indicate that DNA-based hybrid condensates carrying membranous cargo are similarly capable of entering cells, and that the modular targeting and release functionalities demonstrated above could in future be coupled to this capability.

## Conclusions

In summary, we developed hybrid, single and multi-phase biomolecular condensates comprising DNA nanostructures and liposomes, which afford control over the spatial distribution of payload-carrying vesicles and enable dynamic reconfiguration. The distribution of liposomes can be controllably shifted from the surface to the interior of the condensates by changing the density of cholesterol–DNA anchors mediating their interactions with the DNA nanostructures. A simple theoretical model, accounting for the multivalent nature of the liposome-condensate interactions qualitatively reproduces the transition between surface-bound and engulfed states. Orthogonal DNA sequences further enabled vesicle targeting to distinct, co-existing condensates or individual domains within biphasic constructs. Responsiveness was introduced through toehold-mediated strand displacement, enabling sequence-specific release of LUVs that were subsequently captured by separate amphiphilic condensates. The latter functionality is reminiscent of the ability of biological cells to release and re-uptake extracellular vesicles.

This modular platform further demonstrates the potential of nucleic acid–based condensates for building sophisticated biomimetic architectures that imitate both structural and dynamic features of natural membrane-less organelles and other intracellular compartments. While release of LUVs is demonstrated here with condensate disassembly, other strategies such as anchoring LUVs through RNA–DNA hybrids would allow selective release of LUVs by RNase H,^47^ while leaving the DNA network intact.

The design principles demonstrated here, namely those enabling control over liposome engulfment and their targeted release, could be extended to other condensate and coacervate forming materials, such as engineered proteins,^32,34^ RNA^36,37^ and synthetic polymers, ^85^ providing a generalizable strategy for engineering hybrid functional materials. Different materials may offer advantages for biological applications in terms of stability and biocompatibility. DNA, while susceptible to endogenous nucleases, has been found to remain stable in cell culture media supplemented with fetal bovine serum for over 60 hours,^86^ and can be further protected through polymer coatings^87^ or chemically modified nucleotides,^88^ which may also help mitigate immune activation.^89^

By enabling the spatial distribution of liposomes within membrane-less compartments to be programmed, the hybrid condensates will be highly valuable as structural and functional modules in synthetic cell engineering, facilitating spatial organization of biocatalytic pathways achievable by localizing enzymes and/or substrates within the DNA phases, ^46,47^ or encapsulating them in the liposomes. ^90–94^ Vesicle release and re-uptake could then be leveraged for exchanging materials, enabling the construction of synthetic signaling networks. Vesicle exchange between condensate populations could be iterated to build signaling cascades. To this end, one could design DNA-decorated LUVs hosting multiple anchor domains capable of addressing multiple condensates. The LUVs could be immobilized onto, or embedded within, a first condensate population through one anchor type and, upon release, diffuse and localize to a second condensate using another anchor type. The process could then be cascaded multiple times. Targeting would not be restricted to orthogonal base-pairing interactions and could instead rely on protein-binding aptamers, antibodies, or other targeting moieties, enabling interactions with condensates of different chemistry, synthetic cellular systems, and living cells.

Looking ahead to the future deployment of synthetic cellular devices in healthcare, the synthetic extracellular vesicles could serve as vectors for delivering synthetic-cell produced therapeutics to targets in vivo, a direction supported by our proof-of-concept demonstration that hybrid DNA-liposome condensates are internalized by mammalian cells (Fig. S21). Here, additional surface ligands could be introduced for targeted delivery. ^70,95^

On the modeling side, our theory could be further refined to account for liposome deformability and to more accurately evaluate the distribution of available binding sites on the condensate. These refinements, combined with coarse-grained numerical modeling, would provide valuable insights on the biophysical principles governing the surface bound-to-engulfed transition and help draw general design rules applicable to analogous systems.

We expect that our solution for the dynamic organization and triggered release of liposomes from membrane-less compartments will facilitate the design of both internal and external communication pathways in synthetic cellular systems, and between synthetic and biological cells.^96–100^ These capabilities will lower the barriers for deployment of synthetic cell technologies for envisaged applications in biomanufacturing and healthcare.^85,101,102^

## Methods

### DNA Nanostructures Assembly and Handling

DNA nanostructures were designed using NU-PACK^74^ and their correct folding was analyzed using the NUPACK analysis module. All sequences, including the fluorescently-labeled versions, are provided Tables S1-S2, S4-S5.

DNA strands were received freeze-dried and were reconstituted in 1× TE buffer (10 mM Tris, 1 × mM EDTA, pH 8.0). The concentration of reconstituted DNA strands was calculated using the Beer-Lambert Law and the extinction coefficients provided by the manufacturer. Absorbance of reconstituted strands was measured at 260 nm using a Thermo Scientific NanoDrop One UV-Vis spectrophotometer. Reconstituted strands were stored in the fridge at 4°C for short periods of time (< 2 weeks) or in the freezer at - 20°C for longer periods. Before each use, reconstituted strands were vortexed and centrifuged. Stocks of free nanostars (NS A and NS B), linkers (aa, bb, ab) or toeholding linker (aa TH, bb TH) were prepared using the required oligonucleotides and diluted in 300 mM NaCl in 1 × TE to give a final nanostar concentration of 2 µM and final linker concentration of 4 µM. One in five core-forming strands was replaced with its fluorescently labeled counterpart unless otherwise specified (Atto 488 for NS A and Alexa 647 for NS B). The mixtures were then annealed in 0.2 mL 8-tube strips using a thermal cycler (Bio-Rad C1000 Touch Thermal Cycler), by incubating at 95°C for 15 minutes, and then cooling from 90°C to 25°C at a cooling rate of -0.1°C min^−1^.

Nanostructures formed from cholesterol anchors and the docking strand were annealed at a final concentration of 4 µM. The mixtures were annealed in 0.2 mL 8-tube strips using a thermal cycler (Bio-Rad C1000 Touch Thermal Cycler), by incubating at 95°C for 15 minutes, and then cooling from 90°C to 25°C at a cooling rate of -0.1°C min^−1^.

After annealing, stocks of pre-annealed nanostructures were stored in the fridge at +4°C for up to 1 month.

### Preparation of LUVs and Functionalization with DNA Nanostructures

Large unilamellar vesicles (LUVs) were prepared through extrusion at room temperature using an Avanti Research Mini Extruder with 0.1 µm pore size polycarbonate membranes (Whatman Nuclepore).

As-supplied lipids were dissolved in ethanol-stabilized chloroform. A 1 mg lipid film was formed by drying the mixture of lipids (1 mol% Texas Red DHPE, 99 mol% DOPC) under a gentle stream of nitrogen and leaving it under vacuum overnight. The dried lipid film was resuspended at a concentration of 2 mg mL^−1^ in a buffer that matches the osmolarity of the DNA condensates samples (288 mM sucrose and 160 mM NaCl) and vortexed. The hydrated lipid mixture was then subjected to 5 freeze/thaw cycles using liquid nitrogen and a heat block set at 85°C, before being passed through the membrane 21 times. Vesicle mixtures were stored at 4°C in dark microcentrifuge tubes and used within one month of extrusion. Vesicles were then functionalized with DNA nanostructures by overnight incubation at 4°C in the three different ways described below to achieve their localization inside, outside the condensates, or at the interface between the two phases:

### Internally-sequestered LUVs

10 parts cholesterol anchors (4 µM) and 1 part LUVs (as extruded at 2 mg mL^−1^). Then, incubated LUVs were further diluted 5 × in 300 mM NaCl (1 × TE) before adding 0.75 µL to 60 µL DNA samples.

### Surface-tethered LUVs

1 part cholesterol anchors (4 µM) and 1 part LUVs (as extruded at 2 mg mL^−1^). Then, incubated LUVs were further diluted 5 × in 300 mM NaCl (1 × TE) before adding 0.75 µL to 60 µL DNA samples.

### LUVs as linkers

5 parts cholesterol anchors type A (4 µM), 5 parts cholesterol anchors type B (4 µM), and 1 part LUVs (as extruded at 2 mg mL^−1^). Then, incubated LUVs were further diluted 5 × in 300 mM NaCl (1 × TE) before adding to DNA samples in amounts corresponding to the different fractions of ab linker test (*F*_ab_ = {0, 0.05, 0.10, 0.15, 0.20, 0.25, 0.50}).

### Annealing of condensates in 384-Well Plates

DNA condensates were annealed in 384-well plates (IBIDI, µ-Plate 384 Well Glass Bottom # 1.5 Coverslip, sterilized).

Samples were made using pre-annealed stocks of nanostars (2 µM) and linkers (4 µM) and mixed in the defined fractions of ab linker *F*_ab_. For non-toeholding condensates, final nanostars and linkers concentrations were 0.5 µM and 1 µM, respectively. For toeholding condensates, final nanostars and linkers concentrations were 0.75 µM and 1.5 µM, respectively. 20 µL were loaded per well and samples were prepared in triplicate.

If applicable, 1 µL vesicles prepared as described above were added to each well containing 60 µL DNA sample.

We note the modified version of the linker was only used for the phase that was going to be disassembled. For example, in condensates with *F*_ab_ = 0.05 and surface-tethered LUVs on the A-rich phase, the toehold version of the aa linker was used along with the standard version of linker bb.

The annealing protocol included an incubation step at 55°C for 30 minutes to ensure the 6-nt long sticky ends were melted while preserving hybridization in nanostars and linkers, followed by an equilibration step at 40°C for 2 hours. A slow cooling ramp from 40°C to 30°C was then applied at a rate of -0.1°C per 10 min. Finally, samples were cooled to 20°C and stored at +4°C until imaging.

### Delivery of LUVs to Amphiphilic Condensates

C-star amphiphilic condensates were made using the sequences specified in Tables S4-S5, following the protocol described in detail in previous work by Malouf et al. ^45^,^103^. Briefly, C-star condensates were annealed with an initial hold at 95°C for 30 minutes, cooled from 85°C to 50°C at -0.04°C min^−1^, then cooled from 50°C to room temperature at -0.5°C min^−1^. Then 5 µL C-star condensates were added to the wells in which condensates doped with vesicles were annealed, aiming for a final C-star concentration of 0.2 µM. In comparison, [NS A] = [NS B] = 0.4 µM after the addition of C-stars condensates. After initial imaging, the sequence-specific trigger strand (*α* trig. or *β* trig.) was added in a 6 × excess of the toehold strand to the desired well. Disassembly was imaged as described above using epifluorescence microscopy. Confocal micrographs were recorded as end points.

### Confocal Laser Scanning Microscopy for Sample Characterization

Confocal images were acquired with a Leica TCS SP5 microscope using a 40 × /0.85 NA HCX PLAN APO dry objective (Leica). Samples were imaged directly in the 384-well plates in which samples were annealed through a # 1.5 coverslip glass bottom. Atto 488 was excited with a 488 Argon laser and emission was measured between 493 and 543 nm. Alexa 647 was excited with a He/Ne 633 laser and emission was measured between 638 nm and 698 nm. Texas Red DHPE was excited with a He/Ne 594 laser and emission was measured between 599 nm and 629 nm. The pinhole was set to 1 AU.

### Epifluorescence Microscopy for Toehold Mediated Strand Displacement Experiments

Timelapses were performed using a Nikon Eclipse Ti2-E inverted microscope with a Perfect Focus System (PFS) equipped with a Plan Apo *λ* 20 × /0.75 NA, WD 1000 µm dry objective (Nikon), a Lumencor SPECTRA X LED engine and a Hamamatsu Orca-Flash4.0v3 camera.

1 µL trigger strand dissolved in 300 mM NaCl in 1 × TE was added to each well with DNA condensates to a final concentration of 5 µM equivalent to a 6 × excess.

Image segmentation details are provided in the Supplemental Information.

## Acknowledgement

L.D.M., D.A.T., L.M., and C.F. acknowledge support from the European Research Council (ERC) under the Horizon 2020 research and innovation programme (ERC-STG No 851667 - NANOCELL). L.D.M. and C.F. acknowledge support from a Royal Society University Research Fellowship (UF160152, URF \ R \ 221009). R.R.S. acknowledges funding from the Biotechnology and Biological Sciences Research Council through a BBSRC Discovery Fellowship (BB/X010228/1) and from Wolfson College, Cambridge. B.M.M. is supported by a PDR grant of the FRS-FNRS (Grant No. T021024F).

## Supporting Information

The Supporting Information is available free of charge at https://pubs.acs.org/doi/XX.XXXX/acsnano.XXXXXXX.

Supplementary methods, notes, figures, and tables, including oligonucleotide sequences; estimation of anchor coverage (Supplementary Note I); theoretical model of LUV–condensate interactions (Supplementary Note II); LUV characterization by dynamic light scattering; cargo encapsulation and release; radial intensity and partitioning analysis; biphasic condensate morphologies; phase-targeted disassembly and strand-displacement validation; release– capture experiments with amphiphilic condensates; and cellular uptake of hybrid condensates (PDF)

### Associated Content

A version of this manuscript was previously posted as a preprint:

Tanase, D. A.; Malouf, L.; Rubio-Sánchez, R.; Fan, C.; Jain, K.; Mognetti, B. M.; Di Michele, L. Organization and triggered release of liposomes with DNA-based synthetic condensates. 2025, 2025.09.27.678970. bioRxiv. https://doi.org/10.1101/2025.09.27.678970 (accessed June 23, 2026).

### Data Availability Statement

The data supporting the findings of this study are openly available in the University of Cambridge Apollo Repository at https://doi.org/10.17863/CAM.131585.

## TOC Graphic

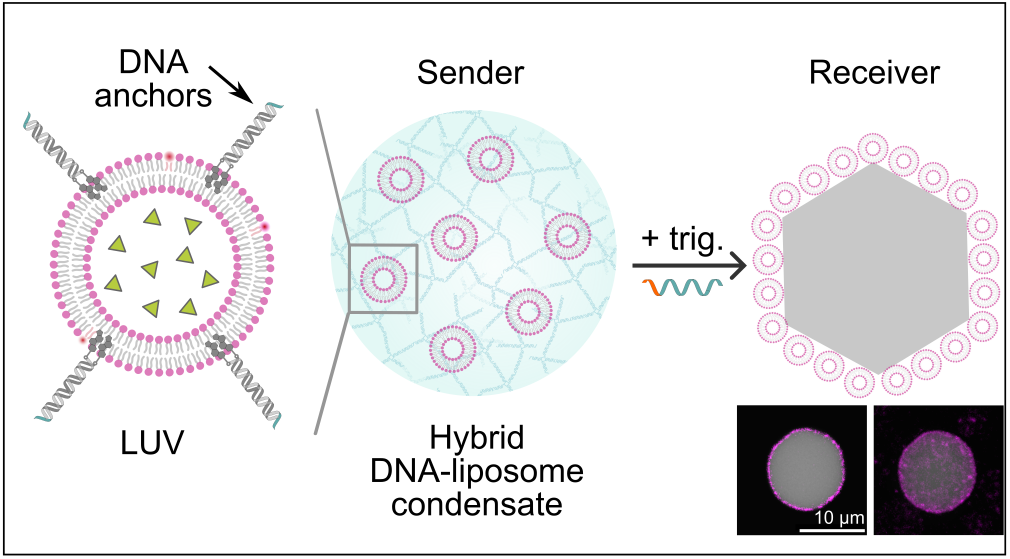

