## Supplementary material for "Organization and triggered release of liposomes with DNA-based synthetic condensates": SI

### Materials

DNA oligonucleotides were purchased from Integrated DNA Technologies (IDT) with the exception of the 5' cholesterol-functionalized strand (Chol_anchor_1) which was purchased from Eurogentec. All strands were purified by the supplier using standard desalting for non-functionalized strands and high-performance liquid chromatography (HPLC) for the purification of cholesterolized strands and fluorophore-labeled strands. 1,2-dioleoyl-sn-glycero-3-phosphocholine (DOPC) was purchased from Avanti Research. 1,2-dihexadecanoyl-sn-glycero-3-phosphoethanolamine, triethylammonium salt (Texas Red DHPE), DNA ladders, BlueJuice Gel Loading Buffer, and SYBR Safe DNA gel stain were purchased from Invitrogen. Sodium chloride, sucrose (RNase- and DNase- free), Sephadex G50, calcein, sodium hydroxide, were purchased from Sigma-Aldrich. 10× TBE buffer and Triton X-100 were purchased from Thermo Fisher Scientific.

### Methods

#### Agarose Gel Electrophoresis

The correct folding of DNA nanostructures along with the implementation of toehold mediated strand displacement was confirmed using agarose gel electrophoresis. Samples were annealed as described above.

Agarose gels were prepared at 1.5% (w/v) in 1 × TBE buffer (89mM Tris, 89mM boric acid, 2mM EDTA, pH of 10 × TBE = 8.3) and stained with 0.01% (v/v) SYBR Safe DNA gel stain. Gels were cast to a thickness of 5 mm and left to set for 30 minutes before covering with 1 × TBE running buffer. Each well was loaded with 12 µL sample composed of ≈ 400 ng DNA and 1.3 × BlueJuice Gel Loading Buffer diluted in 1 × TBE. Equivalent samples were made with a 100 bp DNA ladder (Gene Ruler). Gels were run at a voltage of 80 V (4 V $cm^{-1}$) for 70 minutes and imaged using a Syngene G:Box chemi XX6 system.

#### Vesicle Size Confirmation with DLS

The size of vesicles was confirmed using dynamic light scattering (DLS). DLS backscatter measurements (173°) were performed at 25°C using a Malvern Zetasizer NanoZS equipped with a 633 nm He/Ne laser. After extrusion (and if applicable after incubation with cholesterol anchors), vesicle samples (100 µL diluted to 0.2 mg $mL^{-1}$ lipids) were loaded to a low volume cuvette (ZEN0040).

#### Encapsulation of Calcein in LUVs and Purification

A stock solution of 100 mM calcein was prepared in Milli-Q water, and the pH was adjusted to ≈ 7 using a 1 M NaOH solution.

For experiments confirming cargo retention (Fig. S3), lipid films were rehydrated in a solution of 50 mM calcein dissolved in 300 mM NaCl in 1× TE, and subsequently extruded. Size exclusion chromatography (SEC) was used to separate calcein-loaded vesicles from unencapsulated calcein using a Sephadex G50 resin swollen in a solution of 300 mM NaCl in 1× TE.

For vesicles used for condensates in cell experiments (i.e., incubation with HEK293 cells), a modified buffer system was implemented to preserve vesicle and condensate integrity under physiological conditions. The lipid film was rehydrated in an encapsulation buffer consisting of 50 mM calcein, 235 mM sucrose, and 160 mM NaCl in 1× TE. The sucrose concentration was explicitly reduced by 50 mM relative to the standard rehydration conditions to compensate for the osmotic contribution of calcein.

The corresponding SEC purification was performed using Sephadex G50 resin swollen in an aqueous buffer composed of 285 mM sucrose, 200 mM NaCl, 0.2 mM Tris, and 0.02 mM EDTA in Milli-Q water by incubating for 1 hour at 60°C. Once swollen, the resin was packed under gravity into a 5 mL disposable column (Pierce, Thermo Scientific). A 200 µL aliquot of the extruded LUV mixture was loaded onto the column and eluted using a buffer with an identical composition to the resin-swelling buffer.

Eluted fractions of approximately 200 µL were collected, and the fluorescence intensity of the Texas Red DHPE lipid label was tracked across fractions using a BMG CLARIOstar Plus microplate reader. The peak fluorescent fraction was retained, and the absolute lipid concentration was quantified by interpolating against a standard calibration curve of Texas Red DHPE. Finally, these purified LUVs were functionalized with DNA cholesterol anchors as described previously, accounting for the volumetric dilution introduced during SEC elution to target an average ratio of 125 anchors per vesicle.

#### Fluorimetry to Confirm Cargo Retention

Samples annealed as described above with internally-sequestered LUVs containing encapsulated calcein were used to confirm that the cargo encapsulated in LUVs (i.e., calcein at a self-quenching concentration) is retained inside the vesicles during the one-pot anneal process.

Fluorescence intensity measurements were carried out using a BMG CLARIOstar Plus microplate reader. Since calcein was encapsulated at a self-quenching concentration and the release of calcein is expected to give an increase in fluorescence intensity, the enhanced dynamic range mode was used. Individual measurements consist of 20 flashes per well and were recorded every 20 seconds for 101 cycles ($\approx$30 min).

The optical settings were set as follows:

| Fluorophore | Excitation/nm | Emission/nm | Dichroic filter/nm |
|---|---|---|---|
| Calcein | 480±10 | 530 ±15 | 502.5 |
| Texas Red DHPE | 570±10 | 630±20 | 595 |

2 µL Triton X-100 (0.5 % v/v) was injected in cycle 6 to trigger the lysis of vesicles and the subsequent release of calcein. Each well contained 20 µL hybrid DNA-liposome condensates annealed as described above in "Condensates Annealed in 384-Well Plates".

#### Incubation of DNA-LUV condensates with HEK293 cells

##### Step-wise Buffer Exchange of DNA Condensates for Cell Culture Integration

To mitigate screening-induced structural remodeling or osmotic degradation of the DNA nanostructures upon introduction to cell culture conditions, DNA-LUV hybrid condensates were subjected to a step-wise buffer exchange protocol prior to incubation with cells.

Condensates were initially prepared at a final concentration of 0.75 µM in capillaries using a 300 mM NaCl in 1× TE buffer, functionalized with 0.5 µL of the pre-assembled cholesterol anchor-LUV solution. Three distinct wash buffers were prepared by mixing cell culture media (Dulbecco Modified Eagle's Medium, DMEM, supplemented as described below) with the initial 300 mM NaCl in 1× TE as follows:

- **Wash Buffer 1**: 1 part cell culture media to 2 parts 300 mM NaCl in 1× TE (v/v)

- **Wash Buffer 2**: 2 parts cell culture media to 1 part 300 mM NaCl in 1× TE (v/v)
- **Wash Buffer 3**: 100% cell culture media

The pre-annealed condensates were first extracted into 120 µL of 300 mM NaCl in 1× TE and allowed to sediment under gravity. A 60 µL aliquot of the supernatant was removed and replaced with 60 µL of Wash Buffer 1. The sample was gently agitated to ensure homogeneous mixing and allowed to resediment. This process was iteratively repeated: 60 µL of the supernatant was removed and replaced with 60 µL of Wash Buffer 2, followed by mixing and sedimentation. Subsequently, 60 µL of supernatant was replaced with 60 µL of Wash Buffer 3. Following a final equilibration and sedimentation step, a final 60 µL of supernatant was removed without replacement, yielding isolated, washed condensates for cell delivery.

HEK293 cells (ATCC® CRL-1573™) were grown in Dulbecco Modified Eagle's Medium (DMEM), high glucose, GlutaMAX™ Supplement (ThermoFisher 10566016) containing 10% Fetal Bovine Serum and 1% Antibiotic-Antimycotic (ThermoFisher 15240062) and maintained at 37°C with 5% $CO_2$ in a humidified incubator.

$2.5 \times 10^4$ total cells/mL were seeded the day before the experiment into a 8-well high ibiTreat 1.5 polymer chamber slide with a total volume of supplemented media of 200 µL. On the day of experiment, cell media in each well was aspirated and replaced with 200 µL DMEM with 0.8 µM Hoechst 33342 (Thermofisher 10150888) and cells were incubated at 37°C with 5% $CO_2$ for 10 minutes. The staining media was then aspirated and cells were washed three times with Dulbecco's Phosphate Buffered Saline (Sigma-Aldrich D8537) before a final 200 µL of FluoroBrite™ DMEM (ThermoFisher A1896701) containing 10% Fetal Bovine Serum, 1% Antibiotic-Antimycotic (ThermoFisher 15240062) and 2 mM L-Glutamine (Thermofisher 25030024) was added.

30 µL of the washed condensates were added to the cell samples. Wells were imaged immediately following condensate addition and after a 24 h incubation period during which samples were incubated at 37°C with 5% $CO_2$.

#### Imaging Methods

##### Confocal Laser Scanning Microscopy for HEK293 Cells Characterization

HEK293 cell samples were imaged using a Zeiss LSM 800 confocal microscope, using a Plan-Apochromat 40×/1.3 Oil DIC VIS-IR M27 objective, heated to 37°C with 5% $CO_2$. The fluorescent signal from the Hoechst stain was acquired using a 405 nm laser set to 3.5% intensity, with a collected wavelength range of 400 to 580 nm. The fluorescent signal from the Texas-red labeled lipids was acquired using a 561 nm laser set to 3.0% intensity, with a collected wavelength range 400 to 645 nm. The fluorescent signal from calcein was acquired using a 488 nm laser set to 1.0% intensity, with a collected wavelength range of 400 to 580 nm. Brightfield images were acquired using a ESID Gain of 4. All fluorescent channels were taken with a Master Gain of 500 V and pinhole size of 51 µm. All image frame sizes were set to 512 pixels × 512 pixels with 16 bits per pixel. Line-averaging was enabled and set to 4×. The scan mode along the x direction was selected to be bidirectional. The scanning rate was set to produce a scanning time of 2.06 µs per pixel.

**Image Segmentation** Image segmentation was performed in `MATLAB`[1] (R2024a) using custom-written code and the Image Processing Toolbox.

#### Image Segmentation for Radial Intensity Profiles

Image analysis was implemented to quantify the radial intensity profiles of LUV signal (per condensate) with respect to condensate centroids identified from the condensate channel in Fig. 2**c**.

##### 1. Reference Mask Generation

- Smoothing: A Gaussian filter (`imgaussfilt`) was applied to reduce noise and create smoother object boundaries.
- Background Flattening: To correct for uneven illumination and background signal, a morphological opening was applied using a large, disk-shaped structuring element (`imopen`). This operation removes smaller foreground features while retaining only the slowly varying background intensity, which was then subtracted from the image. The background-subtracted images were adjusted for contrast (`imadjust`).
- Thresholding: Global thresholds were determined using multi-level Otsu thresholding for condensates (`multithresh`).
- Binary masks were created for condensates via image quantization (`imquantize`).

##### 2. Radial Intensity Profile per Condensate with Dilated Mask

- Condensate masks were evaluated for circularity as radial intensity calculations assume a circular shape. A sub-set of $\approx 75\%$ condensates were further analyzed based on the circularity criteria implemented (0.975-1.1).
- Each condensate mask was dilated by 5 pixels to include surface-adjacent signal from LUVs that are surface-bound.
- For each condensate, the average pixel intensity in the LUV and DNA channels was calculated as the sum of pixel intensities divided by the number of pixels.
- Radial bins were defined at 1-pixel intervals up to the condensate radius (dilated by 5 pixels). Then, average intensity was calculated at each radial distance ($I_{\text{LUV}}$) and divided by the average intensity of the condensate ($I_{\text{cond}}$).
- The radial intensity profile of LUV signal was then plotted as a function of radius ($r$) divided by the condensate radius ($R_{\text{cond}}$) and averaged to plot the mean $\pm$ standard deviation.

#### Image Segmentation for LUV Engulfment

Image analysis used to determine LUV localization in Fig. 2**d** for condensates shown in Fig. S4 was designed to quantify how LUVs distribute between the surface and the core of individual condensates as a function of the number of anchors per LUV. The workflow proceeded as follows:

For each condensate, two regions were defined from the segmentation mask: a thin surface shell and an interior core.

- **Surface shell:** An 8-pixel-wide shell was generated at the condensate boundary by dilating the reference mask by 4 pixels and eroding it by 4 pixels, and retaining the region between the two contours.

- **Core:** The condensate core was defined as a concentric region with a radius equal to $0.3\times$ the condensate radius.
- **Depth-corrected intensity ratio:** In both the surface shell and the core, the LUV signal $I_{\text{LUV}}$ was divided by the DNA signal $I_{\text{DNA}}$ measured in the same region. This normalization accounts for the fact that, in a confocal image, light is collected from a slice of finite thickness, so the LUV signal would appear brighter in the core than at the surface even if the LUVs were uniformly distributed.

#### Image Segmentation for 2D Correlation Coefficients

Image analysis used to determine the 2D correlation coefficients reported in Fig. 3**d** and Fig. 4**d** was designed to identify A-rich and B-rich domains from their respective fluorescence channels and compute their correlation with the LUV signal. The workflow proceeded as follows:

1. **Reference Mask Generation (as above)**
2. **Correlation Analysis with LUV Signal**
   - **2D Correlation Coefficients in Fig. 3d endpoint:** Masks were generated per condensate and a median value was computed based on the raw images. It should be noted that masks generated independently from the A and B channels can overlap if pixels are identified as belonging to both the A-rich and the B-rich phase due to segmentation uncertainty. When the two masks overlap only partially ($F_{\text{ab}} < 0.25$), the overlapping region is discarded and only the pixels unique to each mask are retained, so that $r_{\text{A}}$ and $r_{\text{B}}$ report LUV association with A-rich and B-rich regions (see example in Fig. 2**c**).
   - **2D Correlation Coefficients in Fig. 4d timelapse:** For each frame from the second frame onward, the LUV channel (594 nm) was background-subtracted by removing the intensity profile of the final frame in the sequence to account for LUV signal originating from vesicles adhered to the bottom surface of the imaging well. Negative values after subtraction were set to zero. 2D correlation coefficients were then computed. Masks for the A-rich (488 channel) and B-rich (647 channel) domains were generated from a single reference frame and applied consistently to all subsequent frames of the timelapse as the targeted phase was disassembled by the corresponding trigger strand.

#### Image Segmentation and Object Tracking for Fluorescence Quantification of LUVs Deposited on Amphiphilic Condensates

Image analysis for determining the fluorescence intensity measurements reported in Fig. 5**d** and Fig. S19 was designed to identify and track individual or clusters of amphiphilic condensates over time, measuring the fluorescence of the Texas Red DHPE labeled LUVs within each tracked object (i.e., amphiphilic condensates).

##### 1. ROI Initialization and Segmentation

A set of reference Regions of Interest (ROIs) was established by segmenting a single frame after the addition of the trigger strand. The segmentation was performed on the channel corresponding to the amphiphilic C-star condensates (labeled with either Cy5 or fluorescein).

- Image Smoothing: A Gaussian filter (`imgaussfilt`) was applied to reduce noise and create smoother object boundaries.
- Binarization: The smoothed image was binarized using an automatic global threshold (`imbinarize`) to create a binary mask separating objects from the background.
- Mask Refinement: The binary mask was refined by filling internal holes (`imfill`) and subsequently removing small, unwanted objects with an area less than 50 pixels or $\approx 1.3\,\mu m^2$ (`bwareaopen`).
- Initial ROI Definition: Each distinct object in the refined mask was labeled using `bwlabel`, and the centroids of these initial ROIs were calculated and stored as the reference set for tracking.

##### 2. Object Tracking by Centroid Matching

To maintain the identity of each ROI over time, a frame-by-frame tracking algorithm was executed. For each subsequent frame in the timelapse, the following steps were performed:

- Segmentation of current frame: Each new frame was independently segmented using the same procedure described in Section 2 (smoothing, binarization, and refinement) to identify all objects present at that timepoint.
- Centroid-based matching: A correspondence between the ROIs from the previous frame and the objects in the current frame was established using a nearest-neighbor algorithm. The Euclidean distance was calculated between the centroids of the reference ROIs and the centroids of all objects detected in the current frame (`pdist2`). An object in the current frame was considered a match for a reference ROI if the distance between their centroids was less than a predefined threshold (`maxCentroidDist` = 30 pixels $\approx 5\,\mu m$).
- Tracker position update: If a unique match was found, the reference centroid's position was updated to the location of its matched object in the current frame. This iterative updating allows tracking objects as they move across the field of view. If an ROI from the previous frame had no valid match in the current frame (i.e., no object within the `maxCentroidDist` radius), its data for that timepoint was recorded as `NaN` (Not a Number), and its last known centroid position was carried forward for matching in the next frame.

##### 3. Fluorescence and Area Measurement

For each successfully tracked ROI at each timepoint, quantitative measurements were extracted using the object's mask from the current frame.

- Measurement Mask Generation: The binary mask of the tracked object was dilated using a disk-shaped structuring element (`strel`) with a radius of 10 pixels to create a slightly larger measurement area that would include signal from LUVs.
- Fluorescence Quantification: The mean fluorescence intensity of LUVs was calculated from the pixels within this dilated mask in the measurement channel (Texas Red channel) as shown in Fig. S18. Fluorescence intensity was then normalized by the surface area of each ROI. For visualization in Fig. S19, fluorescence trajectories were normalized to the maximum value of their own trace.

### Supplementary Note I: Estimation of Number of Cholesterol Anchors per Vesicle

Below, we provide a rough estimation of number of double cholesterol anchors per vesicle. In the calculations, we make the following assumptions:

1. All vesicles are 100 nm in diameter. Although the membrane filter pore size used for extrusion is 100 nm, our DLS data (Fig. S1) indicates a broader size distribution for the vesicles, meaning they are not all 100 nm in diameter.

2. All lipids used to make the lipid film end up in vesicles. This is very likely an overestimate, since some lipids are likely to be lost during extrusion.

3. All double-cholesterol anchors incubated with vesicles insert in the lipid bilayer. Again, this is likely an overestimate as cholesterolised constructs can stick to the microcentrifuge tube walls.

4. Vesicles are made of pure DOPC. While we generally dope vesicles with Texas Red DHPE, this is added in very small proportions (1 mol%).

#### Estimating the number of vesicles in a DOPC suspension

To estimate the number of unilamellar vesicles in a 2 mg/mL suspension of 1,2-dioleoyl-sn-glycero-3-phosphocholine (DOPC), we assumed all lipid molecules are assembled into spherical unilamellar vesicles of uniform size. The calculation is based on the total number of lipid molecules available and the number of lipids required to form a single vesicle bilayer.

The following parameters were used:

- Lipid concentration: 2 mg/mL
- Molecular weight of DOPC: 786.113 g/mol
- Headgroup area,[2] $a$: $72.5\,\text{Å}^2 = 7.25 \times 10^{-19}\,\text{m}^2$
- Bilayer thickness,[3] $h$: $46\,\text{Å} = 4.6 \times 10^{-9}\,\text{m}$
- Vesicle radius (outer surface), $r$: $50\,\text{nm} = 5.0 \times 10^{-8}\,\text{m}$
- Avogadro's number, $N_\text{A}$: $6.022 \times 10^{23}\,\text{mol}^{-1}$

##### Total number of lipid molecules per µL

We calculate the number of moles of lipids ($n_\text{lipid}$) and number of molecules ($N_\text{lipid}$):

$$n_\text{lipid} = \frac{2\,\text{mg/mL}}{786.113\,\text{g/mol}} = 2.54 \times 10^{-6}\,\text{mol/mL} = 2.54 \times 10^{-9}\,\text{mol/µL}$$

$$N_\text{lipid} = n_\text{lipid} \times N_\text{A} = 2.54 \times 10^{-9} \times 6.022 \times 10^{23} \approx 1.53 \times 10^{15}\,\text{lipids/µL}$$

#### Number of lipids per vesicle

Assuming a bilayer of thickness $h$ for a vesicle with radius $r$ and head group area $a$, the number of lipids per vesicle is estimated for the sum of lipids in both the outer and inner leaflets of the vesicle:

$$N_{\text{lipids/vesicle}} = \frac{4\pi}{a}\left[r^2 + (r-h)^2\right]$$

$$= \frac{4\pi}{7.25 \times 10^{-19}}\left[(5.0 \times 10^{-8})^2 + (4.54 \times 10^{-8})^2\right] = \frac{4\pi \times 4.56 \times 10^{-15}}{7.25 \times 10^{-19}} \approx 7.91 \times 10^4$$

**Estimated number of vesicles per µL**

$$N_{\text{vesicles/µL}} = \frac{N_{\text{lipid}}}{N_{\text{lipids/vesicle}}} = \frac{1.53 \times 10^{15}}{7.91 \times 10^4} \approx 1.93 \times 10^{10}$$

The estimated number of DOPC vesicles of diameter 100 nm in a 2 mg/mL suspension is approximately:

$$\boxed{\approx 1.9 \times 10^{10} \text{vesicles/µL}}$$

#### Estimating the number of double cholesterol anchors per µL

To estimate the number of cholesterol-functionalized DNA anchors in solution, we consider a 4 µM stock solution. The number of molecules per microliter is calculated as:

$$n = [Chol\_anchors] \times V = 4 \times 10^{-6}\,\text{mol/L} \times 1 \times 10^{-6}\,\text{L} = 4 \times 10^{-12}\,\text{mol}$$

$$N = n \times N_{\text{A}} = 4 \times 10^{-12} \times 6.022 \times 10^{23} \approx 2.41 \times 10^{12}$$

$$\boxed{\approx 2.4 \times 10^{12} \text{cholesterol anchors per µL}}$$

Based on the calculations above, we can estimate the number of double-cholesterol anchors per vesicle and the density of cholesterol anchors.

- **Internally-sequestered LUVs:** (incubated with cholesterol anchors and diluted as described in Methods) $\approx 1250$ double cholesterol anchors per vesicle with an anchor density $\rho_{chol} \simeq 3.98 \times 10^{16}$ anchors/m$^2$ in the outer leaflet. Assuming a uniform 2D distribution, spacing between cholesterol anchors can be approximated to 5 nm. Cholesterol mol% $\simeq$ 3.1%.
- **Surface-tethered LUVs :** (incubated with cholesterol anchors and diluted as described in Methods) $\approx 125$ double cholesterol anchors per vesicle with an anchor density $\rho_{chol} \simeq 3.98 \times 10^{15}$ anchors/m$^2$ in the outer leaflet. Assuming a uniform 2D distribution, spacing between cholesterol anchors can be approximated to 15.8 nm. Cholesterol mol% $\simeq$ 0.3%.

As estimated above, cholesterol anchors for both internally-sequestered and surface-tethered LUVs are in a regime below the $\simeq$ 40 mol% solubility limit of cholesterol in phosphatidylcholine bilayers.[4]

### Supplementary Note II: Multivalent Theory Accounting for the LUV-Condensate Interactions

An LUV is mapped onto a rigid sphere of radius $R$ carrying $N_\mathrm{a}$ anchors with mobile tethering points. The relative position between the center of the LUV and the condensate surface is defined by $h$ (Fig. SN1**a**). Consequently, the contact region between the LUV and the condensate, at a given $h$, is $S_\mathrm{CR} = 2\pi R\,(R-h)$ (with $-R < h < R$). The condensate surface presents nanostar (NS) arms with unpaired sticky ends (SEs) at a density equal to $\rho$, where we approximate $\rho = 1/20^2\,\mathrm{nm}^{-2}$. At a given $h$, the total number of NS arms with SEs in the contact region is $N_\mathrm{SE}$, with $N_\mathrm{SE} = \rho\, S_{CR}$.

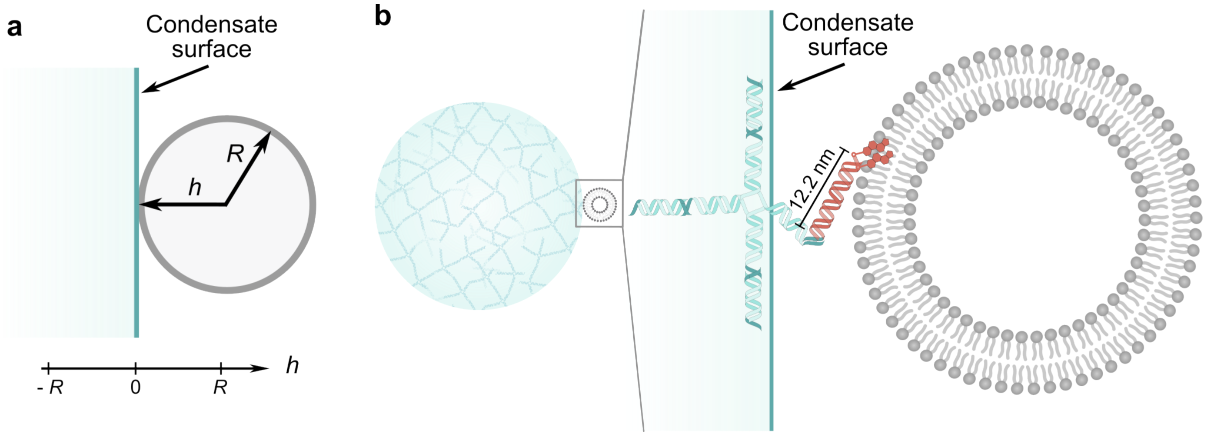


Figure SN1: Schematic showing an LUV interacting with a DNA condensate. **a** The distance between an LUV with radius $R$ and the condensate is defined as $h$. **b** DNA-cholesterol anchors on the LUV surface bind to the sticky ends of NS arms.

We note that, while the model treats LUVs as rigid spheres of radius $R$, vesicles generally have an excess membrane area,[5] allowing them to form finite contact regions without significant condensate deformation. This implies that even at $h = R$, adhesion can occur ($S_\mathrm{CR} > 0$) due to membrane flexibility, whereas our rigid-sphere description underestimates this effect.

**Single bond formation.** Let us consider an anchor constrained to the contact region between the LUV and the condensate, in the presence of a single unpaired NS arm in the contact region (Fig. SN1**b**). The ratio between the probability for the arm and anchor to be bound, over the probability of finding them unbound, is defined by:[6]

$$\tilde{q} = \frac{p_b^{CR}}{p_u^{CR}} = \frac{e^{-\beta \Delta G_0}}{\rho_0 L S_\mathrm{CR}}\,, \tag{S1}$$

where $\beta$ is $1/k_\beta T$, $\Delta G_0$ is the hybridization free energy of the free complementary sequences in bulk, measured at the standard concentration $\rho_0 = 1\,\mathrm{M}$, and $L$ is the length of the anchors ($L = 12.2\,\mathrm{nm}$ for 36 bp anchors). When considering the binding between a condensate SE in the contact region and an anchor with an unconstrained position, the previous relation generalizes as follows:[6]

$$q = \frac{p_b}{p_u} = \frac{e^{-\beta \Delta G_0}}{\rho_0 L S_\mathrm{total}}\,, \tag{S2}$$

with $S_{\mathrm{total}} = 4\pi R^2$. Equations S1 and S2 assume that the condensate SEs are uniformly distributed within the space spanned by the SEs of the anchors in the contact region. The latter assumption is probably an oversimplification. In particular, given the rigidity of the NSs, it is expected that some free energy cost ($\Delta G_{\mathrm{conf}}$) should be paid to reorient a free arm from the condensate towards the LUV surface. Therefore, in the following, we consider that $q$ and $\tilde{q}$ could be reduced by a factor $\exp(-\beta\Delta G_{\mathrm{conf}})$.

**Adhesion free energy, $F(h)$.** We now compute the adhesion free energy of the system at a given $h$, $F(h)$. The multivalent partition function, $Z(h) = \exp[-\beta F(h)]$, reads as follows:[6]

$$Z(h) = \sum_{n_{\mathrm{B}},n_{\mathrm{a}}} Z_h(n_{\mathrm{B}}, n_{\mathrm{a}}) \equiv e^{-\beta \sum_{n_{\mathrm{B}},n_{\mathrm{a}}} \mathcal{F}_h(n_{\mathrm{B}},n_{\mathrm{a}})} \tag{S3}$$

where $n_{\mathrm{B}}$ and $n_{\mathrm{a}}$ are the number of anchor-condensate bonds and total number of anchors (bound or not) in the contact region, respectively. $Z_h(n_{\mathrm{B}}, n_{\mathrm{a}})$ is defined as follows:[6]

$$Z_h(n_{\mathrm{B}}, n_{\mathrm{a}}) = \left[\binom{N_{\mathrm{a}}}{n_{\mathrm{a}}} \frac{(S_{TOT} - S_{CR})^{N_{\mathrm{a}}-n_{\mathrm{a}}} S_{CR}^{n_{\mathrm{a}}}}{S_{TOT}^{N_{\mathrm{a}}}}\right] \left[\binom{n_{\mathrm{a}}}{n_{\mathrm{B}}}\binom{N_{\mathrm{SE}}}{n_{\mathrm{B}}} n_{\mathrm{B}}!\tilde{q}^{n_{\mathrm{B}}}.\right] \tag{S4}$$

$Z_h(n_{\mathrm{B}}, n_{\mathrm{a}})$ comprises a term accounting for the entropic cost of confining $n_{\mathrm{a}}$ anchors in the contact region, along with the free energy contribution of forming $n_{\mathrm{B}}$ bonds out of $n_{\mathrm{a}}$ anchors and $N_{\mathrm{SE}}$ condensate SEs. We approximate $Z(h)$ and $F(h)$ using a saddle point solution:

$$F(h) = \mathcal{F}_h(\overline{n}_{\mathrm{B}}(h), \overline{n}_{\mathrm{a}}) \tag{S5}$$

with

$$\frac{\partial}{\partial n_{\mathrm{a}}}\mathcal{F}_h(\overline{n}_{\mathrm{B}}, \overline{n}_{\mathrm{a}}) = \frac{\partial}{\partial n_{\mathrm{B}}}\mathcal{F}_h(\overline{n}_{\mathrm{B}}, \overline{n}_{\mathrm{a}}) = 0 \tag{S6}$$

The previous equations lead to the following chemical equilibrium balance:

$$\frac{\overline{n}_{\mathrm{B}}}{(N_{\mathrm{a}} - \overline{n}_{\mathrm{B}})(N_{\mathrm{SE}} - \overline{n}_{\mathrm{B}})} = q\,, \tag{S7}$$

where we have used one of the two equations in S6 to express $\overline{n}_{\mathrm{a}}$ as a function of $\overline{n}_{\mathrm{B}}$:

$$\frac{\overline{n}_{\mathrm{a}} - \overline{n}_{\mathrm{B}}}{S_{CR}} = \frac{N_{\mathrm{a}} - \overline{n}_{\mathrm{a}}}{S_{TOT} - S_{CR}}\,. \tag{S8}$$

The multivalent free energy then takes the following, expected, expression:[7,8]

$$\beta F(h) = N_{\mathrm{a}} \log\left(1 - \frac{\overline{n}_{\mathrm{B}}}{N_{\mathrm{a}}}\right) + N_{\mathrm{SE}} \log\left(1 - \frac{\overline{n}_{\mathrm{B}}}{N_{\mathrm{SE}}}\right) + \overline{n}_{\mathrm{B}}. \tag{S9}$$

The total LUV-condensate interaction free energy of the system is then $f(h) = F(h) + S_{CR}\gamma$, where $\gamma$ is the condensate surface tension. The equilibrium penetration depth of the LUV, $h_{\mathrm{eq}}$ is then computed by minimizing $f(h)$.

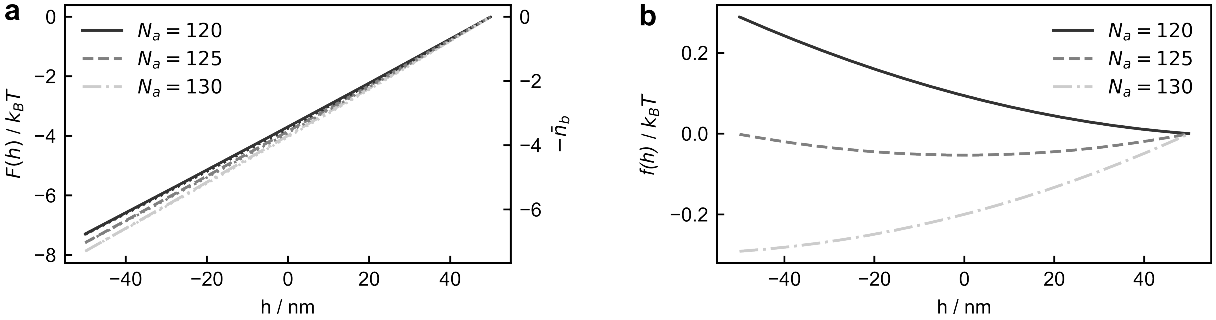


Figure SN2: **The adhesion free energy at the transition point follows a linear regime. a** Multivalent free energy ($F(h)$) and number of bonds (- $n_\mathrm{B}$) for three values of the number of anchors, $N_\mathrm{a}$, inserted in a LUV in the regime in which LUVs are experimentally found to transit from the surface to be internalized in the condensates. In this plot we have $|F(h)/k_\mathrm{B}T + \overline{n}_\mathrm{B}| < 0.6$, validating the approximation used in Eqs. S10. In particular, the number of bonds featured by the system increases linearly from $\overline{n}_\mathrm{B} = 0$ for $h = R$ to $\overline{n}_\mathrm{B} \approx 7$ for $h = -R$. **b** Total free energy of the system ($f(h) = F(h) + S_{CR}\gamma$), where $\gamma$ is the condensate surface tension. In both panels, $\Delta G_0 = -12\,k_\mathrm{B}T$ and $\Delta G_\mathrm{conf} = 6.74\,k_\mathrm{B}T$.

In Figs SN2**a** and SN2**b**, we show $F(h)$ and the total free energy ($f(h)$) close to the transition point between surface-bound ($h_\mathrm{eq} = R$) and fully-embedded ($h_\mathrm{eq} = -R$) states, respectively. $F(h)$ follows a nearly linear trend in $h$ (and therefore in the contact area, $S_{CR}$). It follows that $f(h)$ is nearly linear, resulting in an equilibrium point equal either to $h_\mathrm{eq} = R$ or to $h_\mathrm{eq} = -R$. This observation explains the abrupt transition from surface-bound to fully-embedded states reported in Fig. S5. A linear trend is found in the limit in which the number of bonds $n_\mathrm{B}$ is much smaller than $N_\mathrm{SE}$ and $N_\mathrm{a}$. In this limit we have

$$\beta F(h) = -\overline{n}_\mathrm{B} \qquad \overline{n}_\mathrm{B} \approx N_\mathrm{SE} N_\mathrm{a} q \sim S_{CR}. \tag{S10}$$

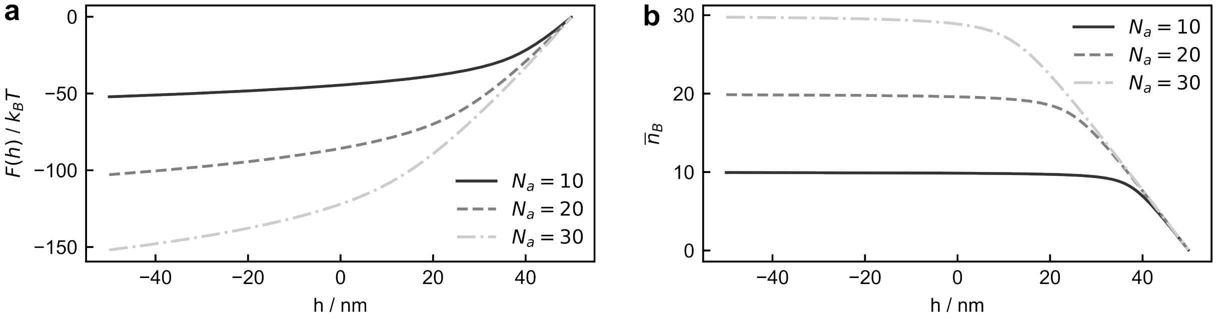


Figure SN3: **Non-linear adhesion free energies emerge under bond saturation conditions. a** Adhesion free energy, $F(h)$, and **b** number of bonds, $\overline{n}_\mathrm{B}$, in saturation conditions. The low number of anchors, $N_\mathrm{a}$, results in SEs on NS arms being saturated. In both panels $\Delta G_0 = -20\,k_\mathrm{B}T$ and $\Delta G_\mathrm{conf} = 6.74\,k_\mathrm{B}T$. Saturation conditions also require stronger bonds than what was used in Fig. SN2.

Nonlinearities in $F(h)$ become relevant when going towards the saturation regime in which the number of bonds is limited either by the number of condensate NS arms with available SEs or by the number of anchors, $\overline{n}_\mathrm{B} \approx \min[N_\mathrm{a}, N_\mathrm{SE}]$. For example, in Fig. SN3 we use $N_\mathrm{a} = 10$, which causes the anchors to become readily saturated. In this regime, increasing the number of NS SEs no longer provides a proportional decrease in free energy.

**Accounting for polydispersity in LUV size and anchor density.** We study the effect of the fluctuations in the size of the LUVs and the number of anchors on the penetration transition

(Fig. S5). We fit the LUV radius distribution obtained in DLS experiments (Fig. S1) using a log-normal distribution

$$R \; = \exp\left(\mu + W\sigma\right) \qquad W \sim \mathcal{N}(0,1) \tag{S11}$$

where $\mu = 4.01$, $\sigma = 0.35$, and $W$ is sampled from a normal distribution. Similarly, we model the number of anchors ($N_a^{\mathrm{fluc}}$) using a Poisson distribution with average equal to $N_a$ (the nominal value of the number of anchors)

$$P(N_a^{\mathrm{fluc}}) = \frac{1}{N_a^{\mathrm{fluc}}!}(N_a)^{N_a^{\mathrm{fluc}}} e^{-N_a} \approx_{N_a \gg 1} \frac{1}{\sqrt{2\pi N_a}} \exp\left(-\frac{(N_a^{\mathrm{fluc}} - N_a)^2}{2N_a}\right) \tag{S12}$$

where, in the large $N_a$ limit, we have approximated $P$ with a gaussian distribution.

The theoretical results of Fig. 2**d** ($p_{\mathrm{theory}}$) are then obtained as follows: For each value of $N_a$, we sample 4000 independent size and number of anchors realizations, $\{R_\alpha, N_{a,\alpha}^{\mathrm{fluc}}\}_{\alpha\in[1,4000]}$, using Eqs. S11 and S12. For each LUV realization, we calculate the equilibrium penetration depth $h_{\mathrm{eq},\alpha}$ following the procedure detailed above. A LUV is considered engulfed when $h_{\mathrm{eq},\alpha}$ is smaller than a threshold value $h_{\mathrm{thr}}$, which is chosen equal to $h_{\mathrm{thr}} = -0.75 \cdot R$. If $h_{\mathrm{eq},\alpha} > h_{\mathrm{thr}}$, LUV $\alpha$ is said to be at the surface. Finally, $p_{\mathrm{theory}}$ is calculated as the ratio between the number of LUVs at the surface and the number of engulfed LUVs. The specific value of $h_{\mathrm{thr}}$ has little impact on $p_{\mathrm{theory}}$ given the sharp transition observed in monodisperse systems (Fig. S5).

### Supplementary Figures

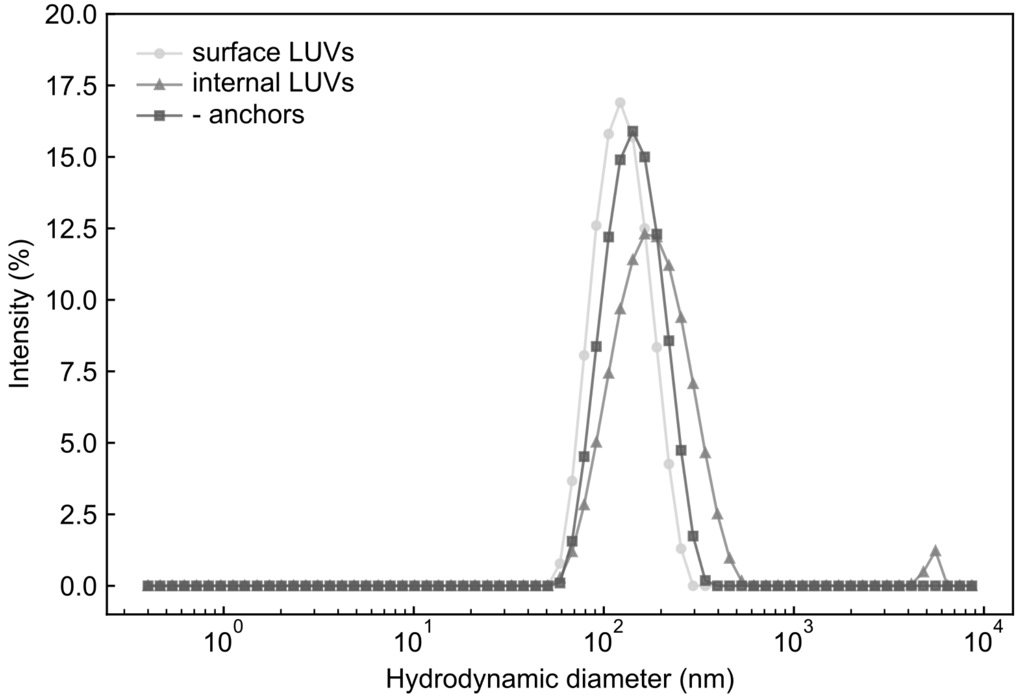


Figure S1: **Dynamic light scattering (DLS) confirms the hydrodynamic diameter of extruded LUVs.** Dynamic light scattering confirms that the hydrodynamic diameter of LUVs is close to their nominal size (∼100 nm) and slightly increases when including anchors with increased density. Each curve is the average of three measurements and the measurements are shown for anchors with sticky ends complementary to those on nanostars B.

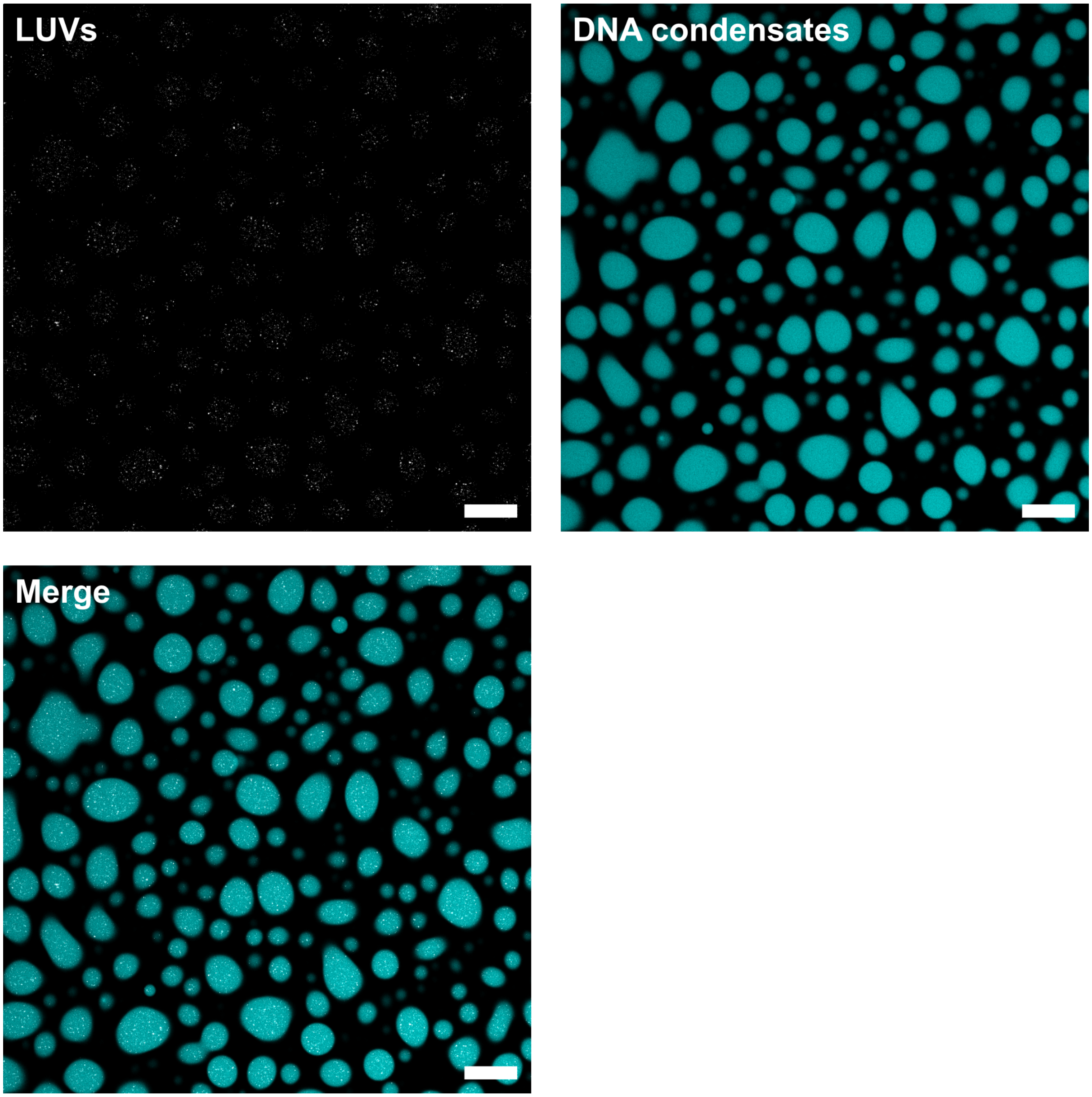


Figure S2: **Large field of view corresponding to the sample highlighted in Fig. 1, showing confocal micrographs of hybrid DNA-liposome condensates.** The LUV channel (TR-DHPE) is scaled from minimum to maximum pixel intensity to aid visualization, while the DNA condensate channel (Alexa 647) is shown without intensity adjustment. Scale bars: 20 µm.

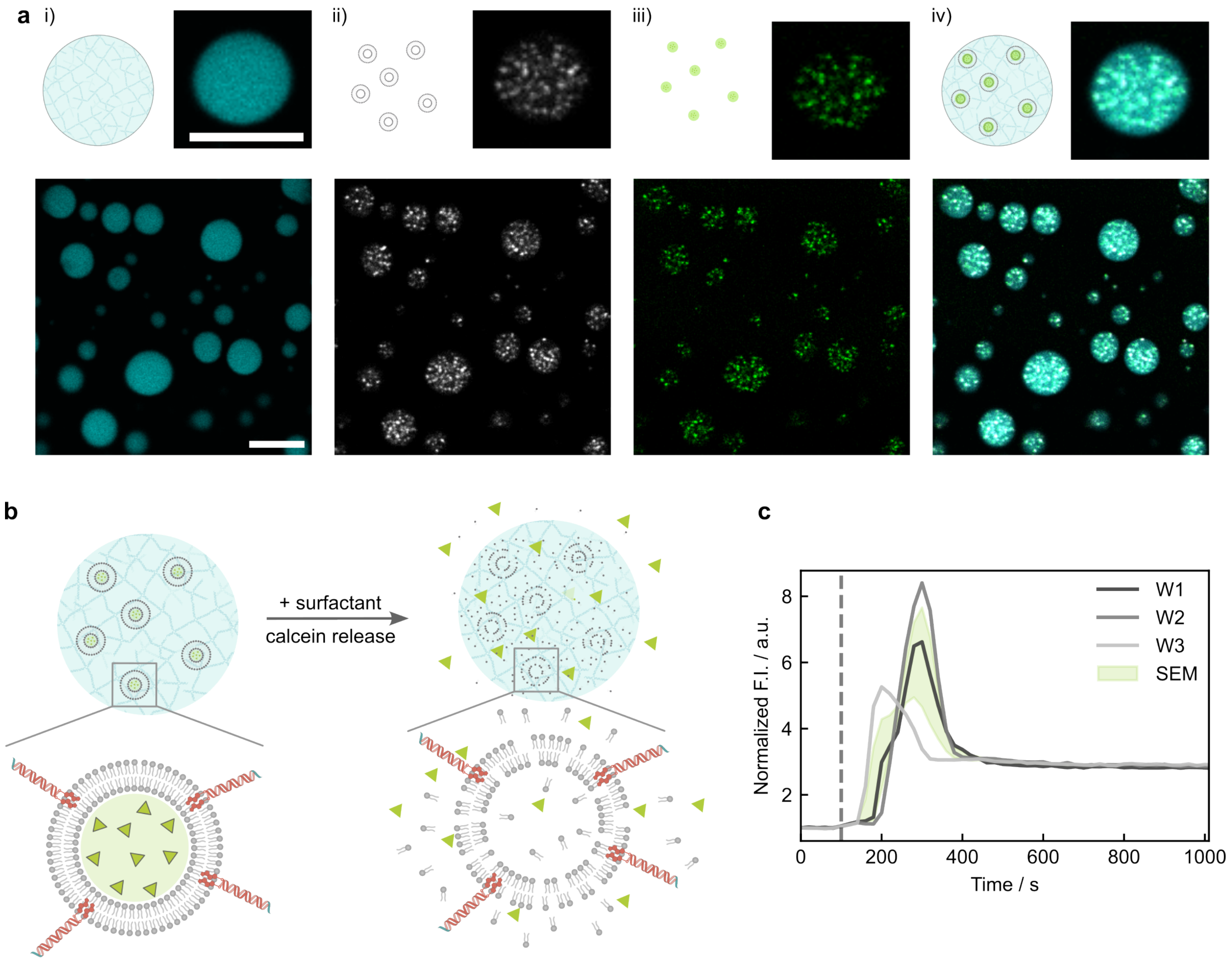


Figure S3: **Cargo (calcein) remains encapsulated in LUVs during the one-pot annealing process that forms hybrid DNA–liposome condensates.** **a** Confocal micrographs and a zoomed-in view of a condensate showing: i) condensate labeled with Alexa 647, ii) vesicles doped with Texas Red DHPE, iii) cargo signal (calcein, detectable by confocal microscopy even at self-quenching concentrations), and iv) overlay of the three channels. Scale bar: 10 µm. **b** Schematic of the calcein release experiment. Addition of the surfactant Triton X-100 lyses vesicles and releases encapsulated calcein, which increases fluorescence due to dilution of the self-quenching concentrations. **c** Calcein fluorescence intensity measured in three wells (W1, W2, W3) during the release assay, normalized to the initial fluorescence. The dashed line indicates the time of surfactant addition. After a short diffusion period, vesicle lysis releases calcein, leading to a surge in fluorescence intensity that subsequently plateaus as calcein diffuses away from the focal plane. The green shaded region indicates the standard error of the mean (SEM) across the three wells. In this sample, we used cholesterol anchors that bind a 5‘ toehold-modified linker (Table S2), mirroring the toehold design later used for toehold-mediated strand displacement (TMSD). In the other experiments, cholesterol anchors are bound via NS SEs, leaving the linker toehold free to drive TMSD reactions.

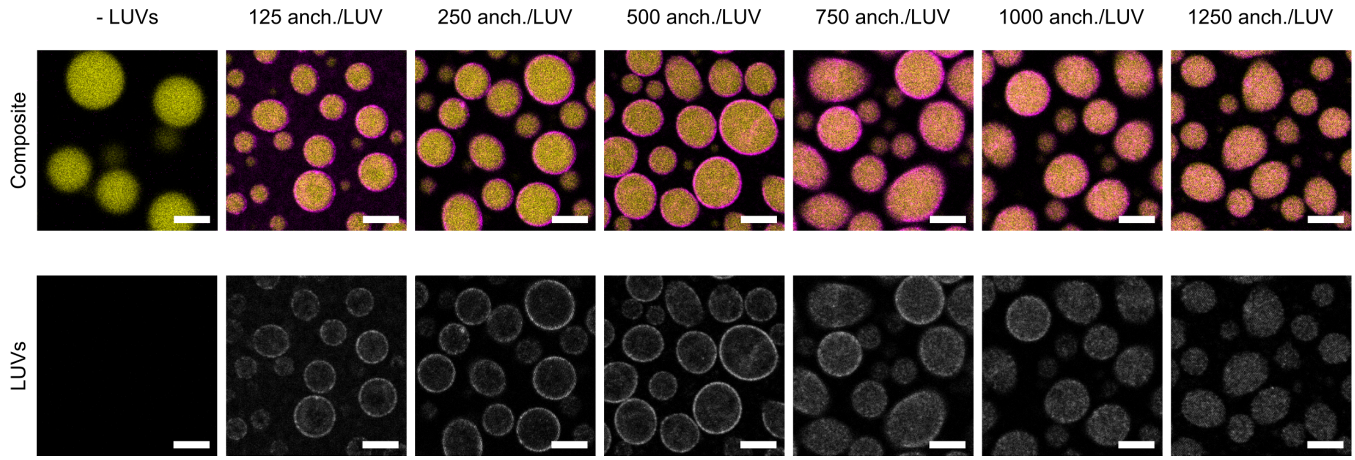


Figure S4: **Confocal micrographs showing hybrid DNA-LUV condensates with varying numbers of cholesterol anchors per LUV.** With 125 anchors/LUV, vesicles are localized on the surface of DNA condensates. Increasing the number of cholesterol anchors per LUV drives the internalization of vesicles into the condensate core. Top: merged channels scaled linearly from 0 to maximum pixel intensity, showing DNA condensates in yellow (labeled with Atto 488) and fluorescently labeled LUVs in magenta (Texas Red DHPE). Bottom: pristine grayscale images of LUV signal. Condensates were made using nanostars A and linkers aa with sequences specified in Table S1. The same dilution of LUVs was used across all samples. Samples were prepared in triplicate, with representative data shown here; the corresponding quantitative analysis is detailed in Fig. **2d**. Scale bars: 10 µm.

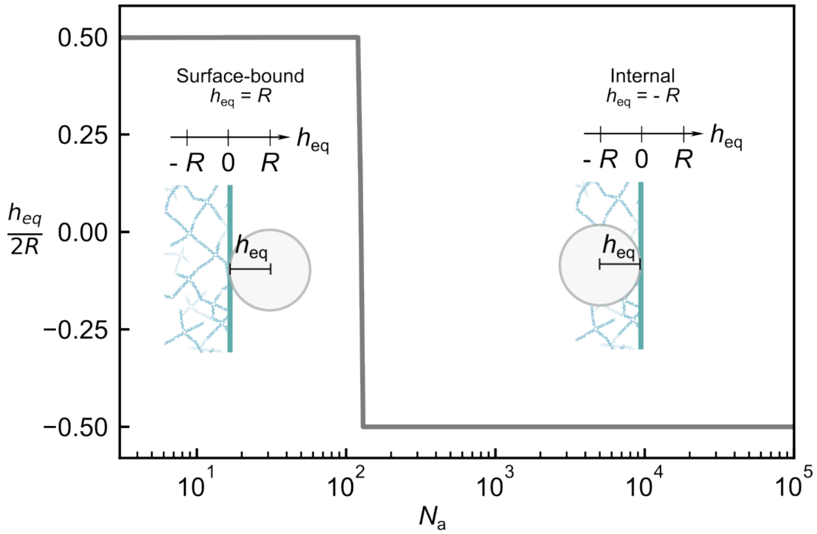


Figure S5: Computed equilibrium penetration depth of an LUV, $h_{eq}$, normalized by the LUV diameter, $2R$, as a function of the number of DNA anchors per vesicle ($N_\mathrm{a}$) when using $\Delta G_\mathrm{conf} = 6.7\, k_B T$. Other model parameters were fixed, as outlined in the main text and Supplementary Note II.

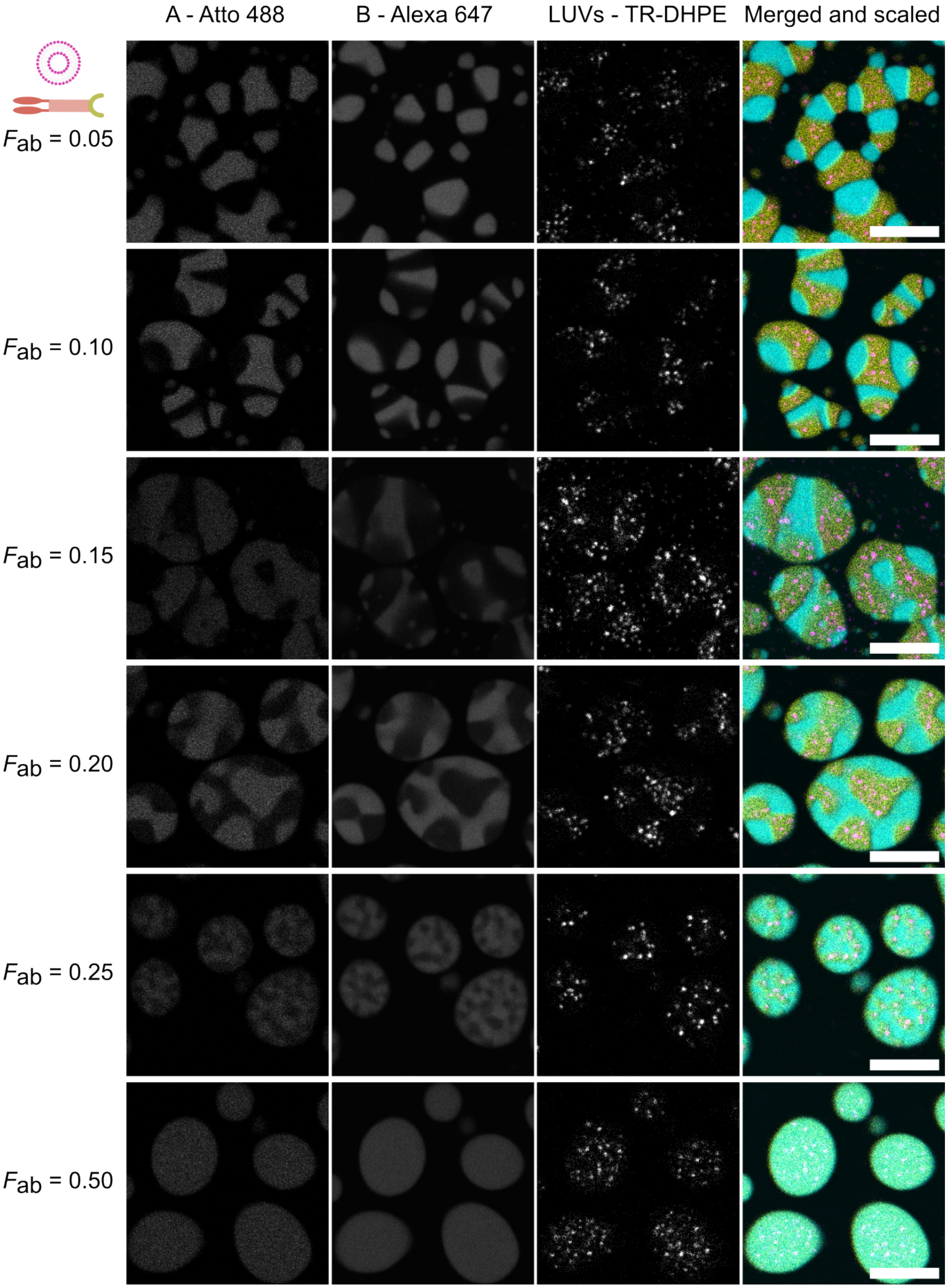


Figure S6: **Individual grayscale images along with the merged image as shown in Fig. 3 for the different fractions of the ab linker, $F_{ab}$, with vesicles internally-sequestered in the A-rich phase (yellow).** The A-rich and B-rich phases are shown unscaled. LUVs labeled with Texas Red DHPE are scaled for better visualization from 0 to maximum pixel intensity. The merged image is shown as in the main text with all channels scaled from 0 to the maximum pixel intensity. Scale bars: 10 µm.

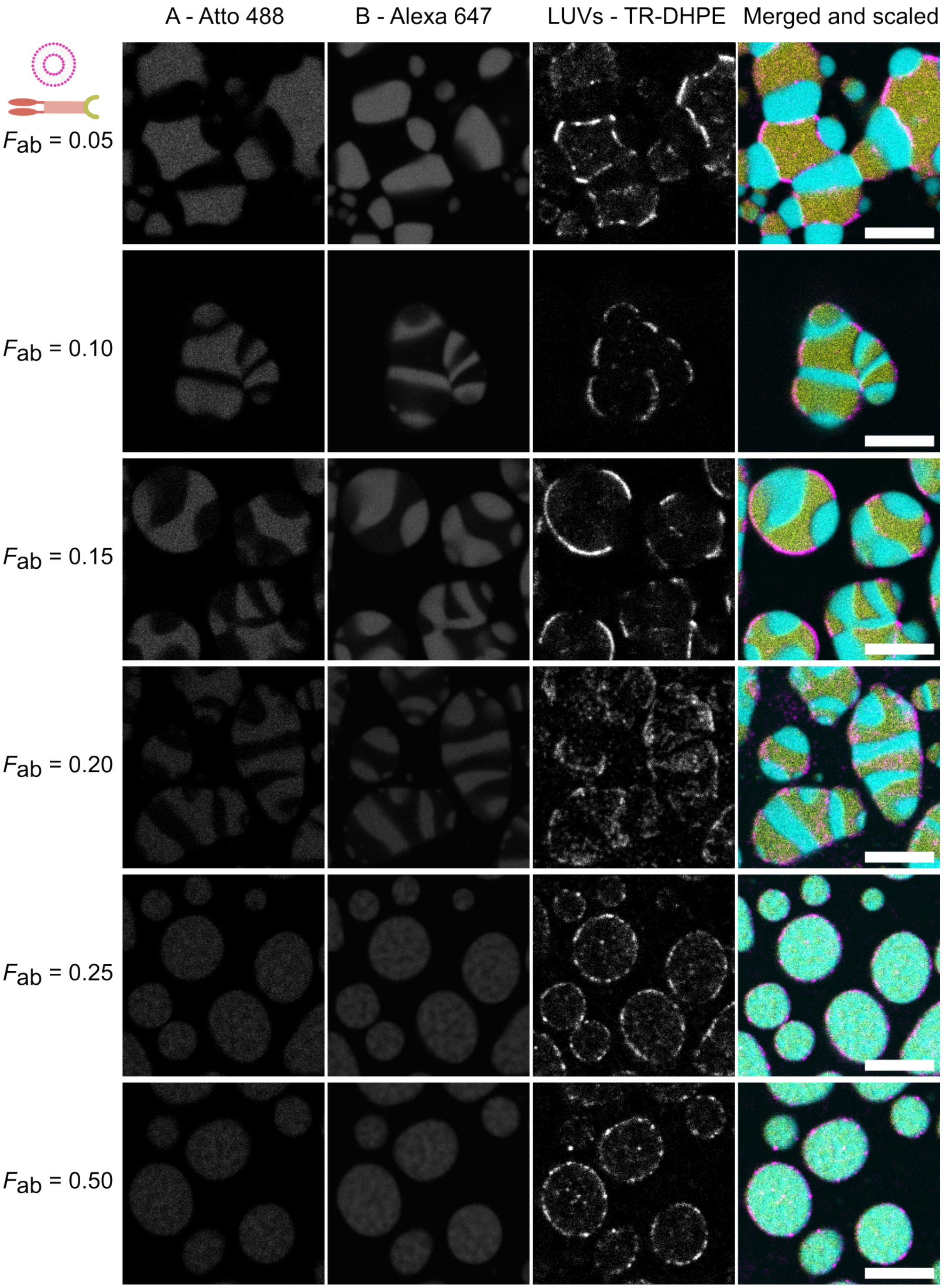


Figure S7: **Individual grayscale images along with the merged image as shown in Fig. 3 for the different fractions of the ab linker $F_{ab}$ with vesicles associated on the surface of the A-rich phase (yellow).** The A-rich and B-rich phases are shown unscaled. LUVs labeled with Texas Red DHPE are scaled for better visualization from 0 to maximum pixel intensity. The merged image is shown as in the main text with all channels scaled from 0 to maximum pixel intensity. Scale bars: 10 μm.

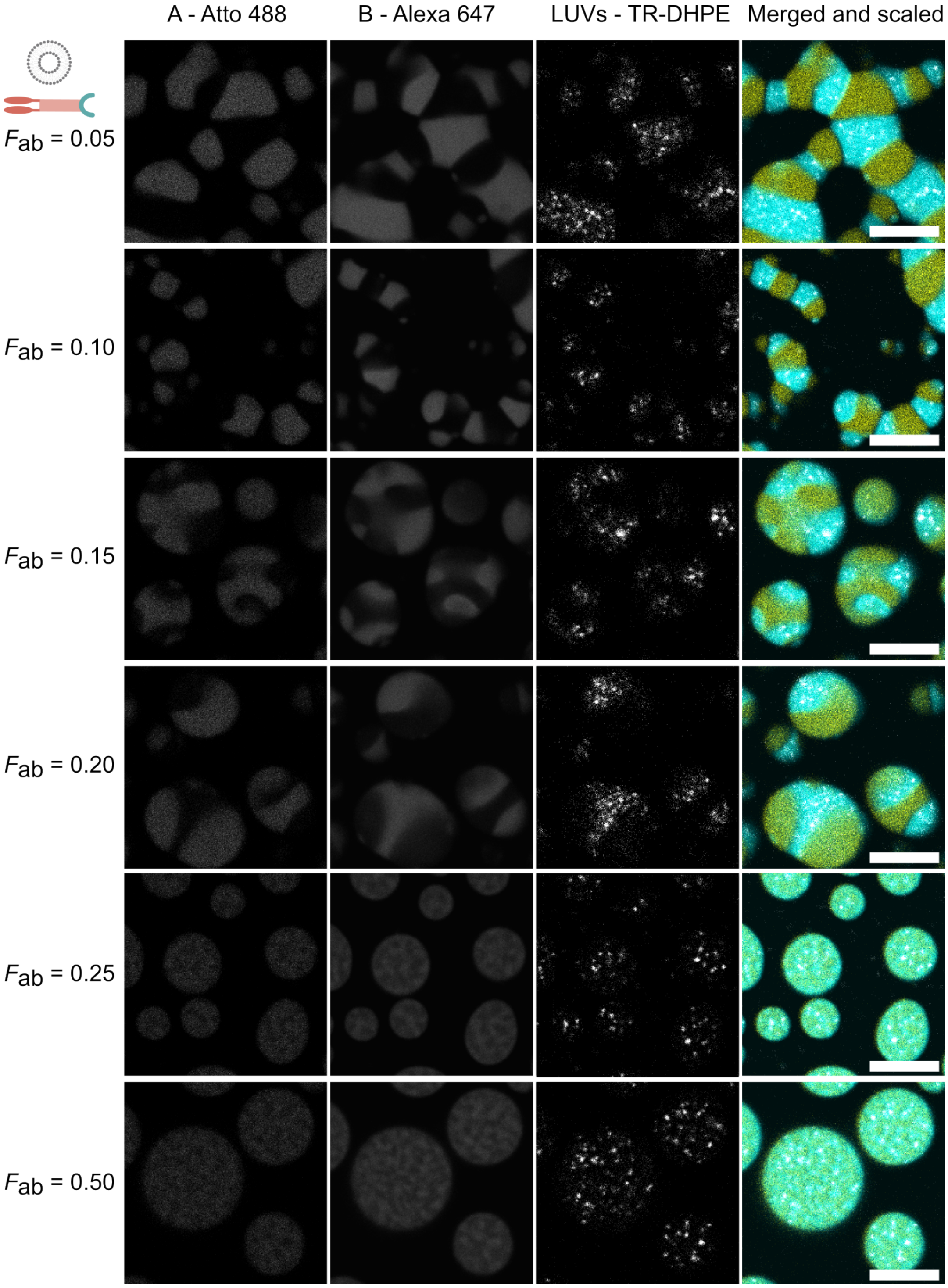


Figure S8: **Individual grayscale images along with the merged image as shown in Fig. 3 for the different fractions of the ab linker, $F_{ab}$, with vesicles internally-sequestered in the B-rich phase (cyan).** The A-rich and B-rich phases are shown unscaled. LUVs labeled with Texas Red DHPE are scaled for better visualization from 0 to maximum pixel intensity. The merged image is shown as in the main text with all channels scaled from 0 to the maximum pixel intensity. Scale bars: 10 µm.

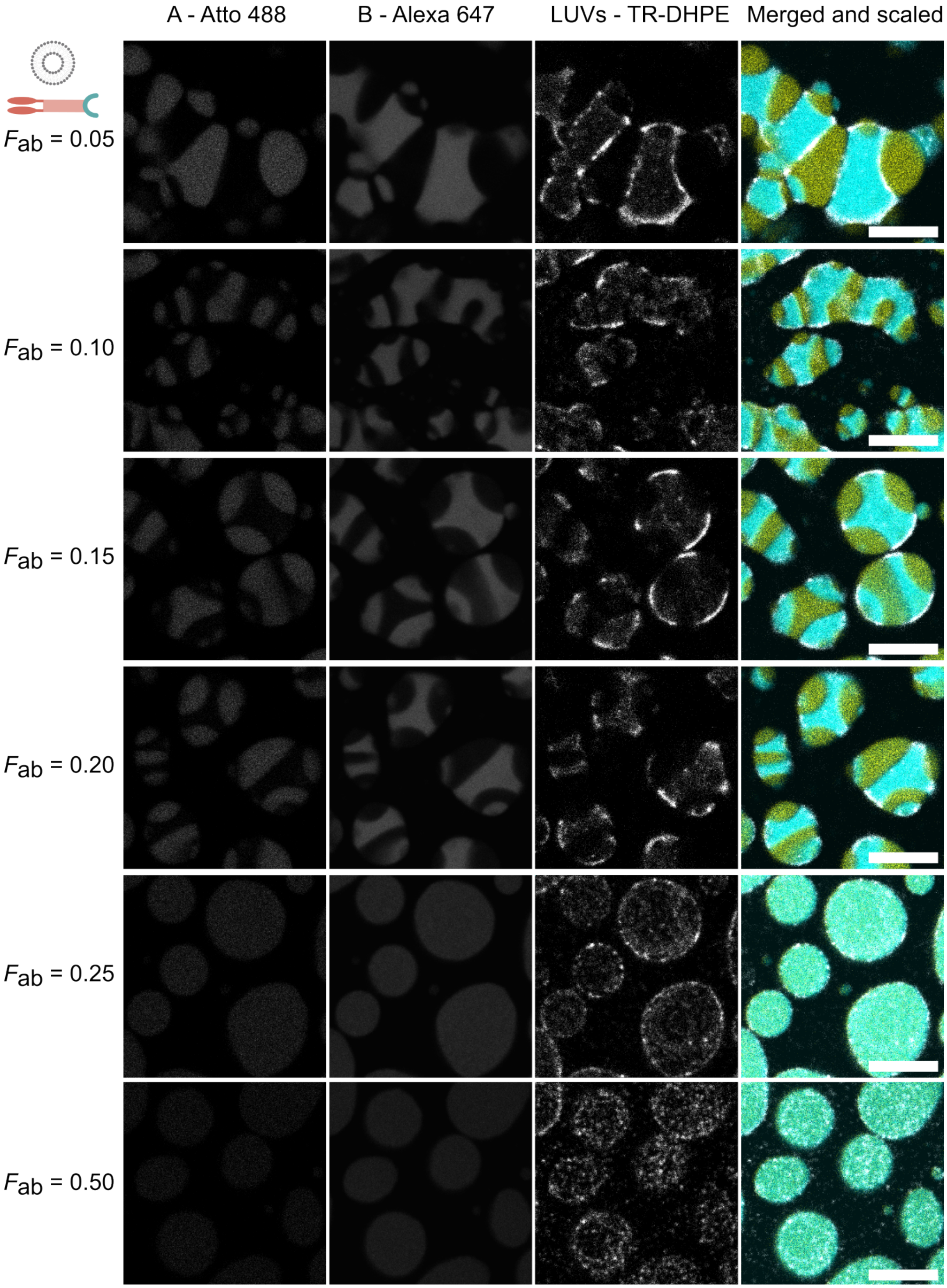


Figure S9: **Individual grayscale images along with the merged image as shown in Fig. 3 for the different fractions of the ab linker, $F_{ab}$, with vesicles associated on the surface of the B-rich phase (cyan).** The A-rich and B-rich phases are shown unscaled. LUVs labeled with Texas Red DHPE are scaled for better visualization from 0 to maximum pixel intensity. The merged image is shown as in the main text with all channels scaled from 0 to the maximum pixel intensity. Scale bars: 10 µm.

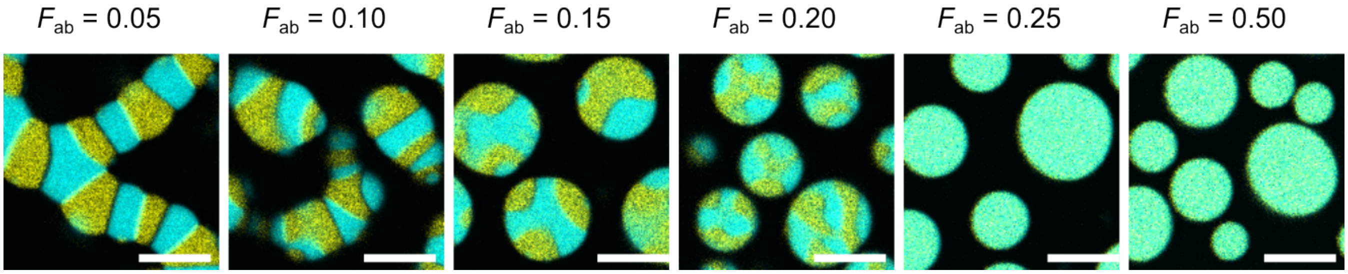


Figure S10: **Negative control showing confocal micrographs of condensates lacking LUVs, with varying fractions of the ab linker, $F_{ab}$.** Condensates with LUVs and different $F_{ab}$ are shown in Fig. 3. Confocal micrographs were scaled from 0 to the maximum pixel intensity. Scale bars: 10 µm.

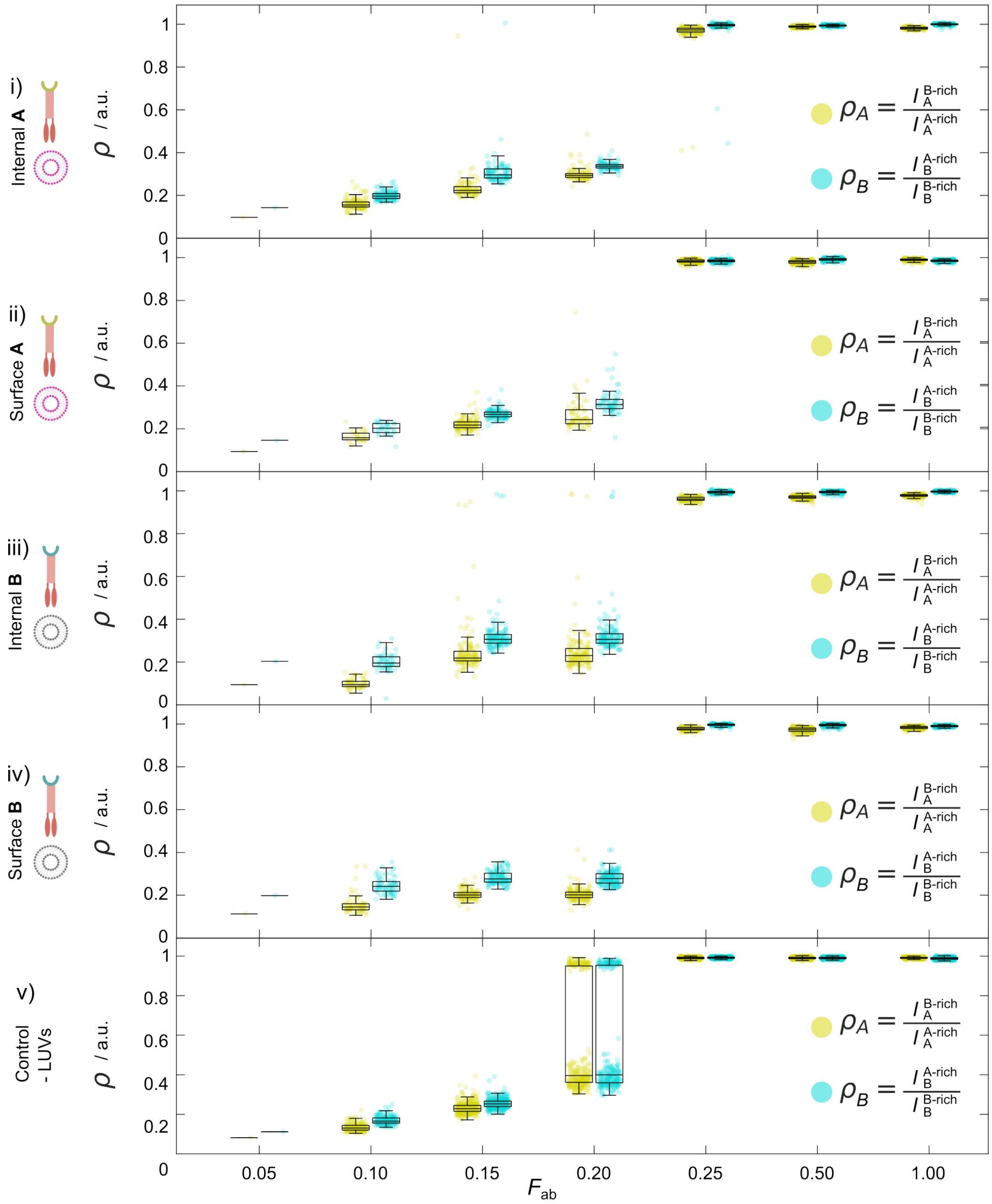


Figure S11: **Partition coefficients ($\rho_A$ and $\rho_B$) for the four types of samples shown in Fig. 3, and Fig. S10 as a function $F_{ab}$, the fraction of ab linker.** To quantify mixing in biphasic condensates, we calculate the partition coefficients, $\rho_A$ and $\rho_B$. Each coefficient is defined as the ratio of fluorescence intensity for a given NS type (A or B) measured in the phase depleted of that NS type (B-rich or A-rich, respectively) to the intensity measured in the corresponding enriched phase (A-rich or B-rich, respectively). Thus, $\rho_A$ and $\rho_B$ approach 0 in fully de-mixed condensates and approach 1 in fully mixed (monophasic) condensates. As mixing increases for higher $F_{ab}$ values, partition coefficients increase. Only one value is reported for $F_{ab} = 0.05$ because condensates formed long-range networks, rather than individual condensates and one value was calculated for the entire network. For $F_{ab} > 0.05$, each data point represents a value calculated per condensate. Box plots show the interquartile range and the median value. Yellow and blue markers denote partition coefficients values for the A-rich and B-rich phase, respectively.

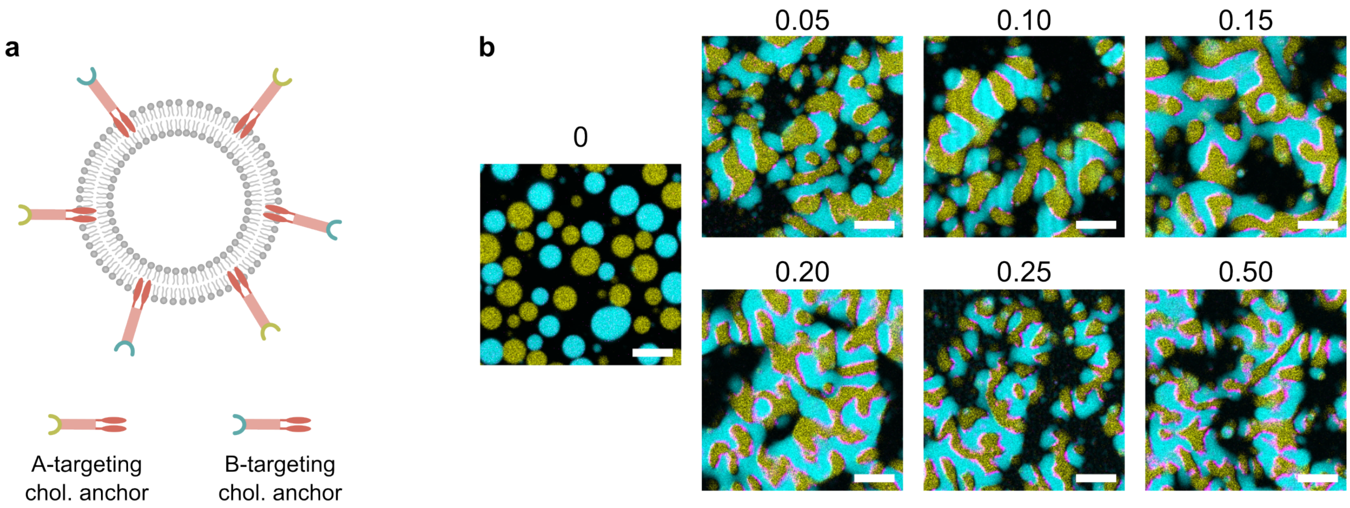


Figure S12: **LUVs incubated with both anchor types serve as linkers between NS A and NS B in the absence of ab DNA linkers.** **a** Schematic of a vesicle incubated with equimolar amounts of cholesterol anchors targeting NS A and NS B, enabling simultaneous binding to both nanostar types and accumulation at the interface. **b** Confocal micrographs (scaled from 0 to maximum pixel intensity in each channel) for different fractions of LUVs incubated with both A- and B-targeting cholesterol anchors in a 1:1 ratio. The indicated fractions correspond to the equivalent fractions of ab linker. Scale bars: 10 µm.

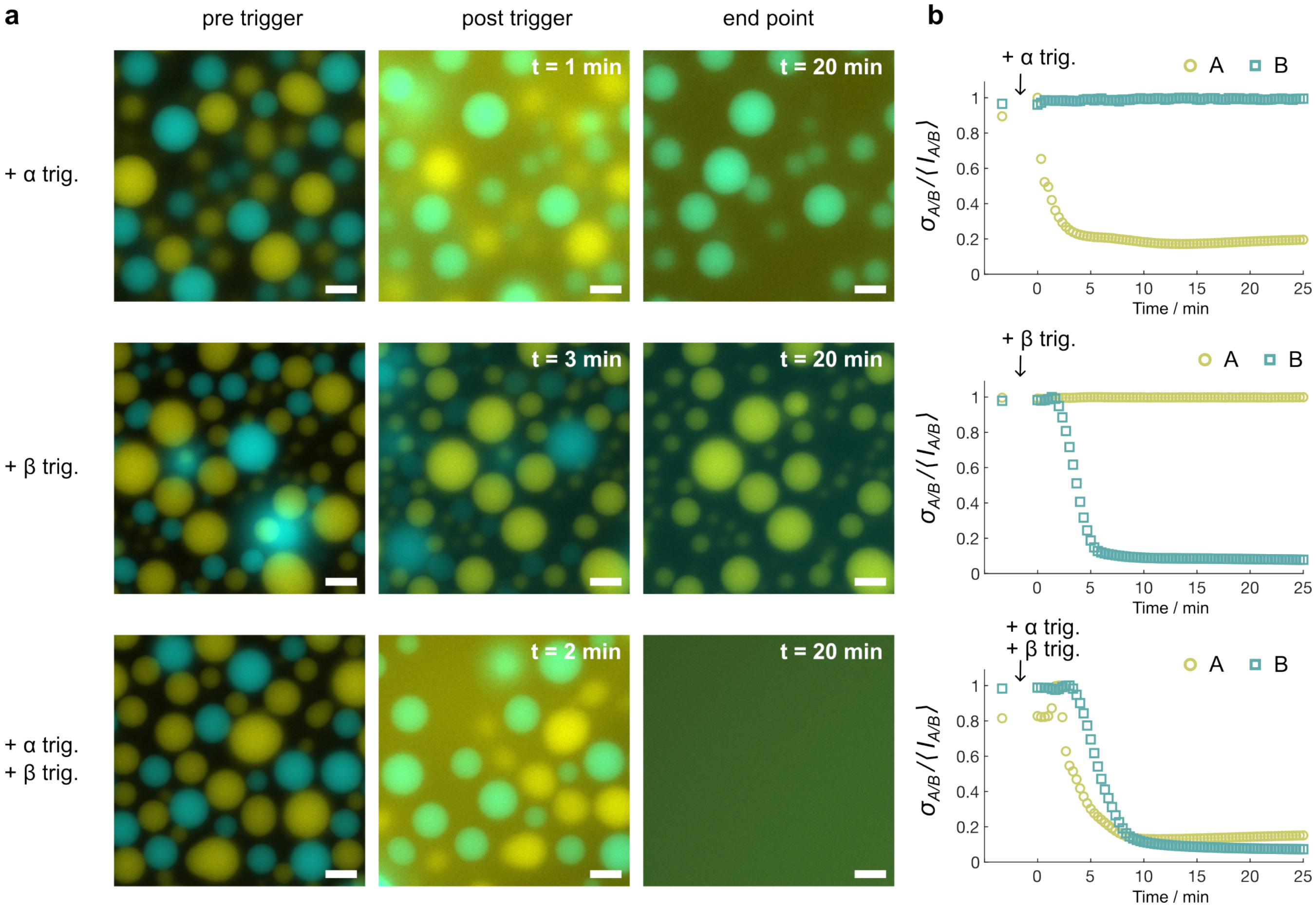


Figure S13: **Epifluorescence micrographs showing targeted disassembly in mixed samples containing orthogonal condensates (A in yellow and B in cyan) at three key time points. a** The addition of the $\alpha$ trigger strand leads to the disassembly of A condensates (yellow, top). The addition of the $\beta$ trigger strand leads to the disassembly of B condensates (cyan, middle), and addition of both triggers leads to disassembly of both A and B condensates. Highlighted post-trigger frames were captured at different time points due to differences in diffusion time of the pipetted trigger strands, which determines the onset of the strand displacement reaction. All micrographs were contrast-adjusted using the minimum pixel intensity and 10% above the maximum intensity in the first (pre-trigger) frame. **b** Normalized coefficient of variation at each time point, for condensates made with NS A (yellow, circle markers) and NS B (cyan, square markers). $\sigma_{\mathrm{A/B}}$ and $\langle I_{\mathrm{A/B}}\rangle$ are the standard deviation and the mean of the pixel values, respectively, evaluated in the full field of view at each time point. A drop in standard deviation indicates condensate disassembly. Scale bars: 10 µm.

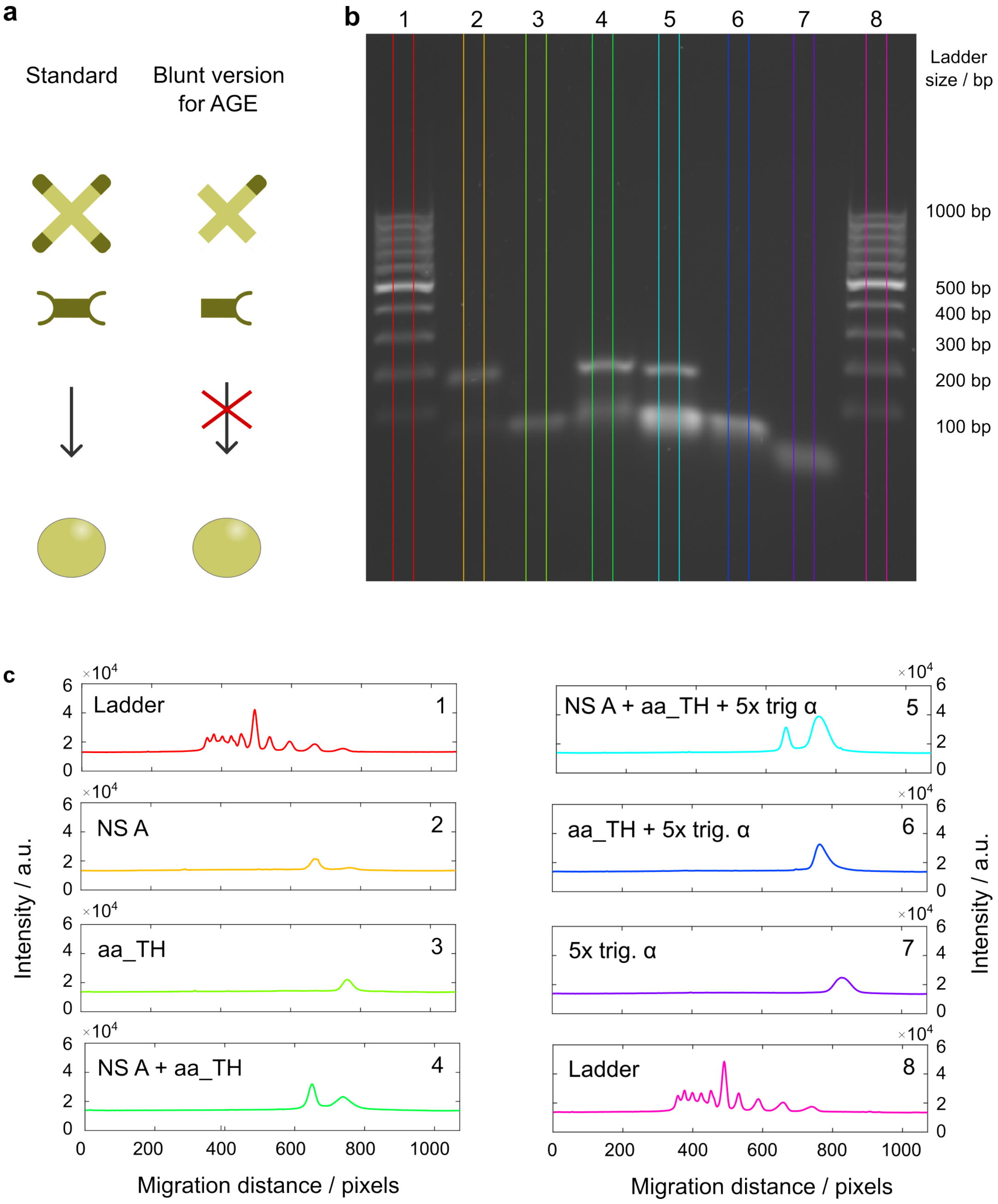


Figure S14: **Agarose gel electrophoresis (AGE) confirms the correct folding of DNA constructs (NS A, linker aa with toehold) and disassembly in the presence of trigger strand $\alpha$.** **a** Schematic showing modified versions of nanostar A and linker aa_TH with blunt ends to prevent the formation of condensates. **b** Image of the agarose gel with a 100 bp ladder (L) on each side (lanes 1 and 8). Lane 2: blunt nanostar A (with only one arm having a sticky-end). Lane 3: linker aa_TH with only one sticky-end . Lane 4: NS A + aa_TH. Lane 5: NS A + aa_TH + 5× excess $\alpha$ trigger. Lane 6: aa_TH + 5× excess $\alpha$ trigger. Lane 7: 5× excess $\alpha$ trigger. **c** Lane intensity profiles for the samples in panel **b**. The sample type and lane number are specified on each plot (top left and right, respectively).

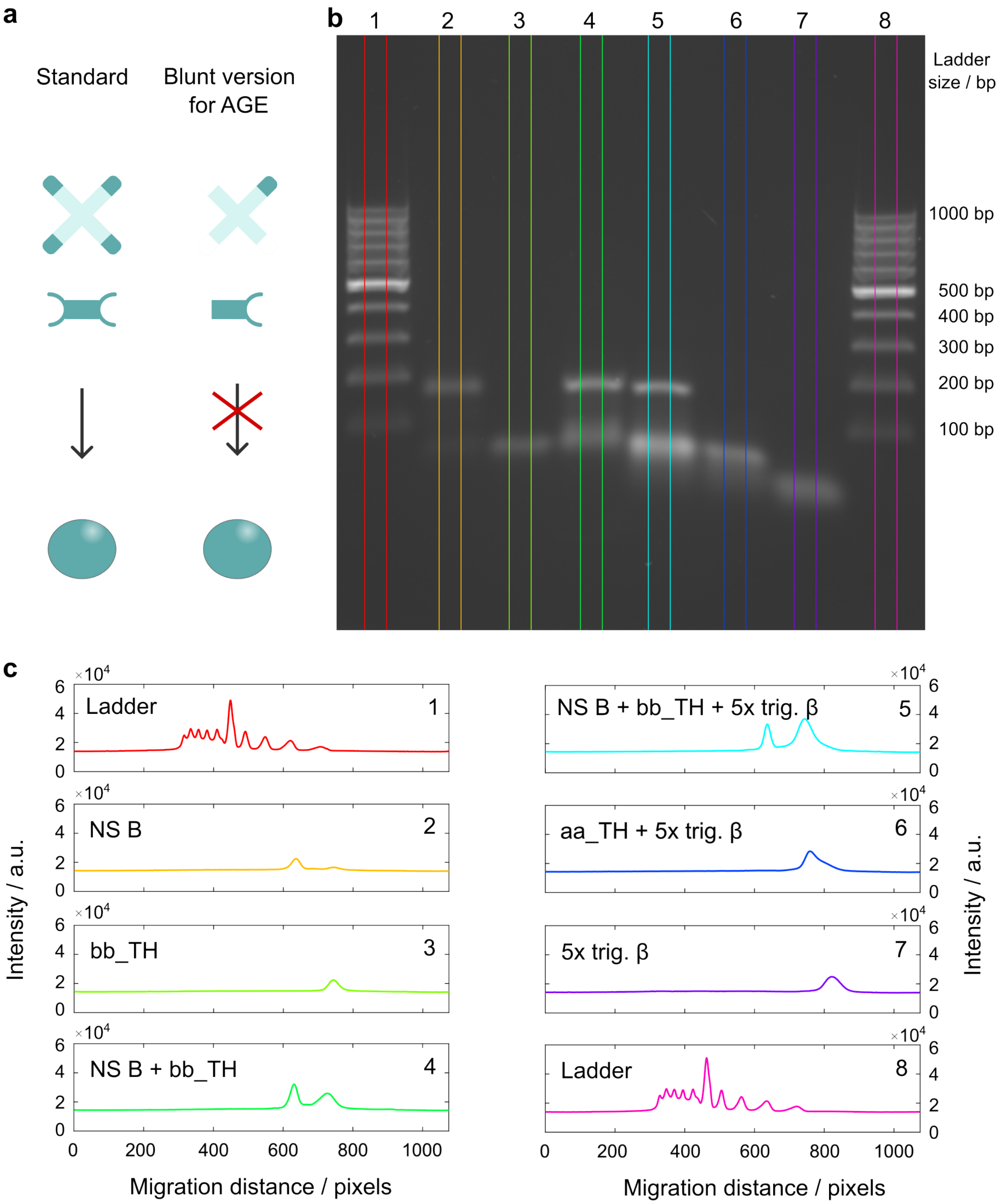


Figure S15: **Agarose gel electrophoresis (AGE) confirms correct folding of DNA constructs (NS B, linker bb with toehold) and disassembly in the presence of trigger strand $\beta$.** **a** Schematic showing modified versions of nanostar B and linker bb_TH with blunt ends to prevent the formation of condensates. **b** Image of the agarose gel with a 100 bp ladder (L) on each side (lanes 1 and 8). Lane 2: blunt nanostar B (with only one arm having a sticky-end). Lane 3: linker bb_TH with only one sticky-end . Lane 4: NS B + bb_TH. Lane 5: NS B + bb_TH + 5× excess $\beta$ trigger. Lane 6: bb_TH + 5× excess $\beta$ trigger. Lane 7: 5× excess $\beta$ trigger. **c** Lane intensity profiles for the samples in panel **b**. The sample type and lane number are specified on each plot (top left and right, respectively).

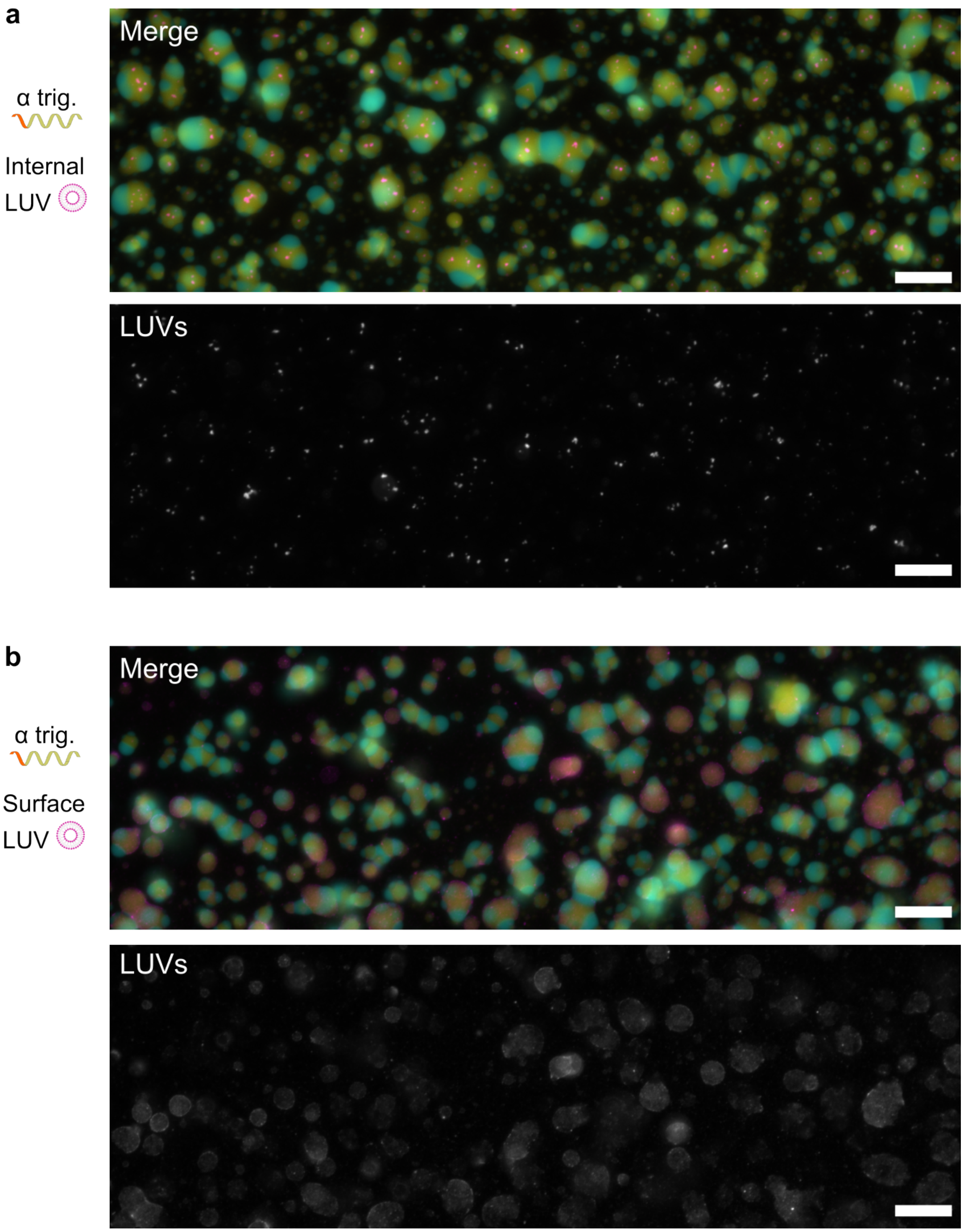


Figure S16: **Large fields of view of epifluorescence micrographs highlighted in Fig. 4b for the sequence-specific release of LUVs from the A-rich phase, before the addition of trigger $\alpha$. a** Internally-sequestered LUVs in the A-rich phase (as in Fig. 4**bi**) shown in merged micrograph (top) and the LUV channel in grayscale (bottom). **b** Surface-tethered LUVs in the A-rich phase (as in Fig. 4**bii**) shown in merged micrograph (top) and the LUV channel in grayscale (bottom). Channels corresponding to the two DNA phases were scaled from minimum pixel intensity to 1.5× the maximum value and the LUVs channel was scaled from minimum to maximum pixel intensity. Scale bars: 20 µm.

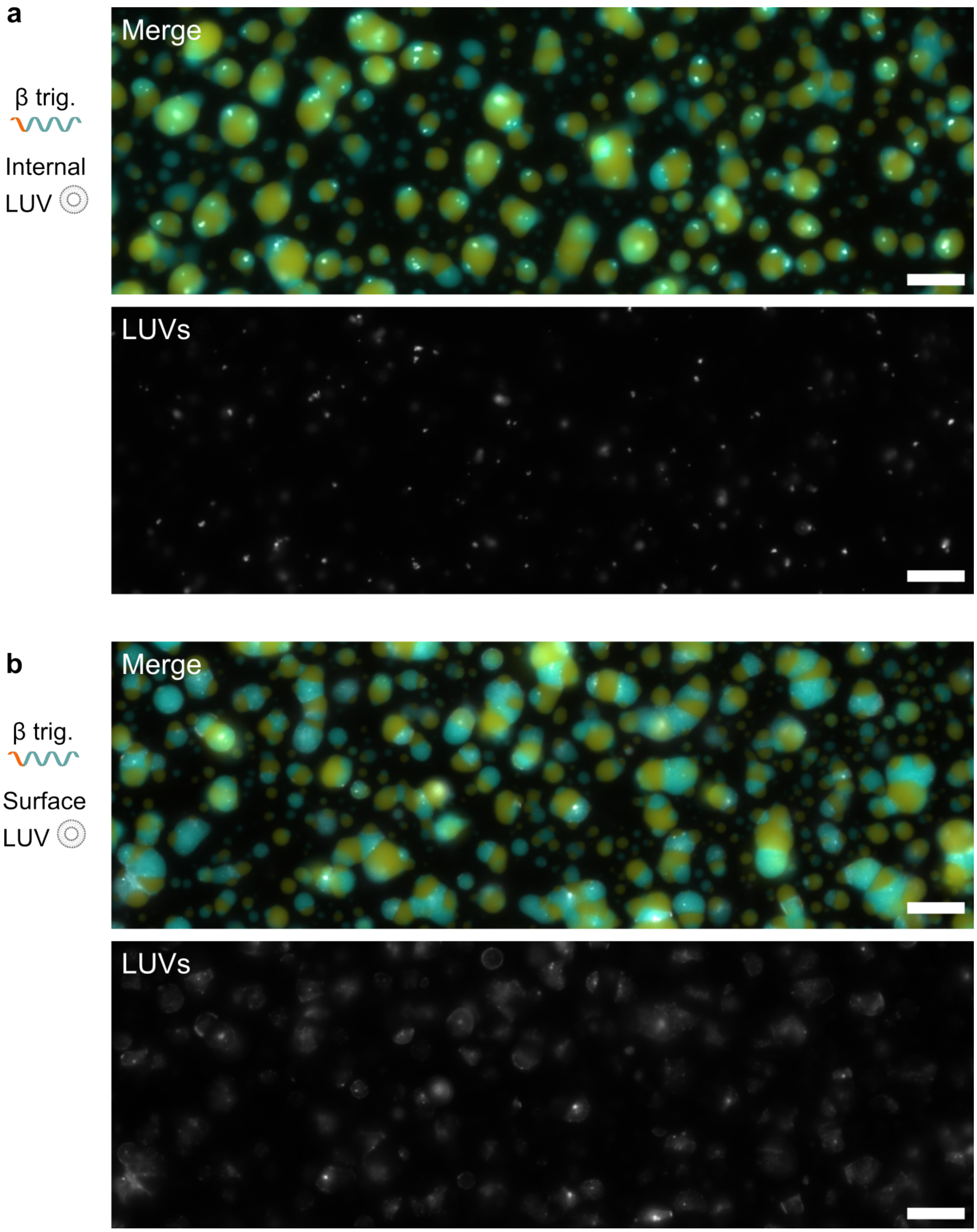


Figure S17: **Large fields of view of epifluorescence micrographs highlighted in Fig. 4b for the sequence-specific release of LUVs from the B-rich phase, before the addition of trigger $\beta$. a** Internally-sequestered LUVs in the B-rich phase (as in Fig. 4**b**iii) shown in merged micrograph (top) and the LUV channel in grayscale (bottom). **b** Surface-tethered LUVs in the B-rich phase (as in Fig. 4**b**iv) shown in merged micrograph (top) and the LUV channel in grayscale (bottom). Channels corresponding to the two DNA phases were scaled from minimum pixel intensity to 1.5× the maximum value and the LUVs channel was scaled from minimum to maximum pixel intensity. Scale bars: 20 µm.

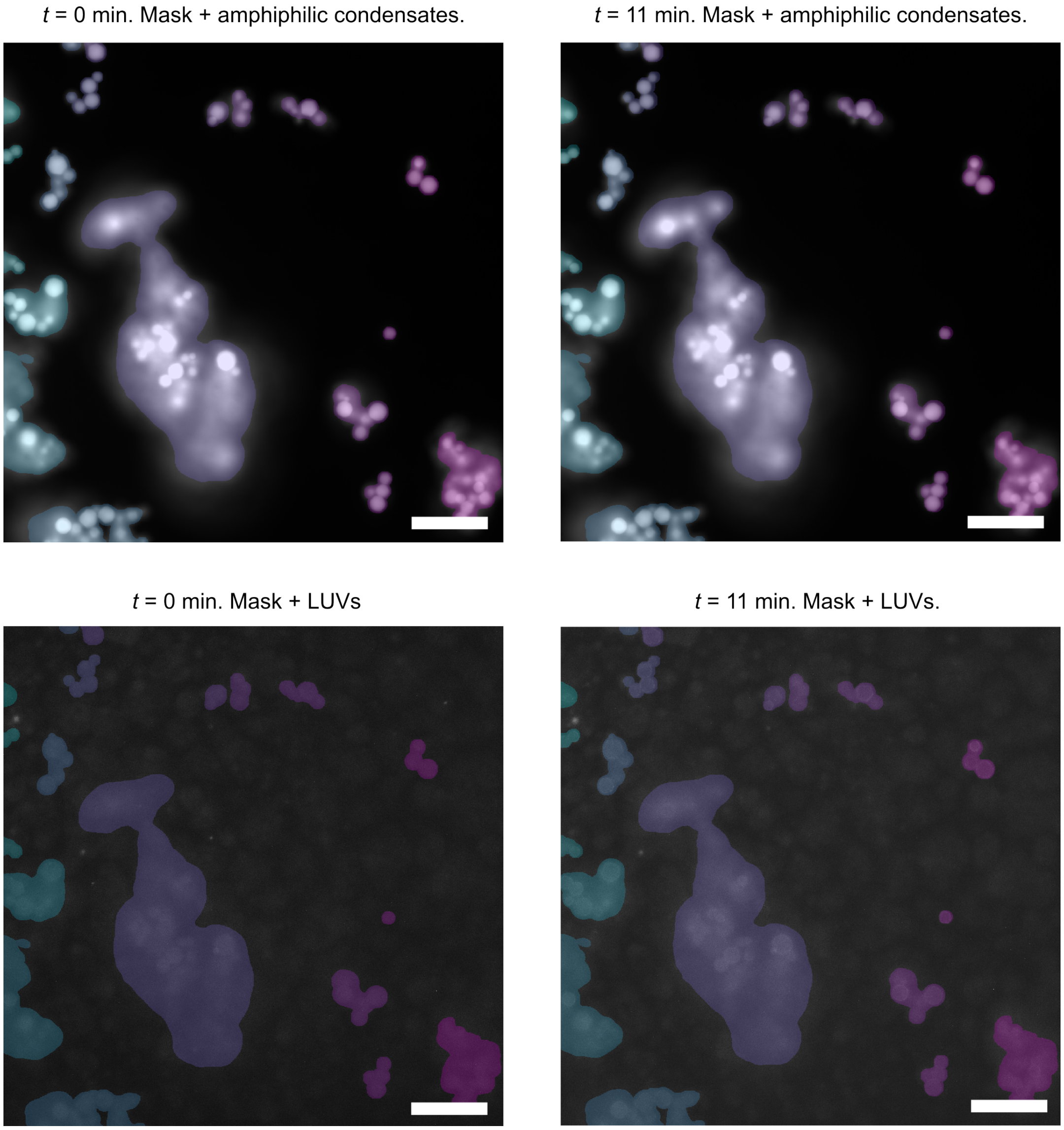


Figure S18: **Epifluorescence micrographs overlaid with masks for monitoring LUV fluorescence intensity over time as surface-tethered LUVs are released from DNA condensates (made with NS A) and deposited onto amphiphilic condensates.** Top: Masks as identified for receiver amphiphilic condensates overlaid with micrographs of the condensates (Cy5) before the release of LUVs (left) and after the release of LUVs (right). Bottom: Same masks as identified for receiver amphiphilic condensates overlaid with the LUV channel before the release of LUVs (left) and after the release of LUVs (right). Different false colors represent different ROIs in which fluorescence is monitored. Note: partial coating with LUVs is noticed even at early stages which could be due to an excess of LUVs in the sample. Masks are identified by thresholding and expanding by 10 pixels (≈1.6 µm) in all directions to ensure signal coming from the outside of the condensates is taken into account. Scale bars: 50 µm.

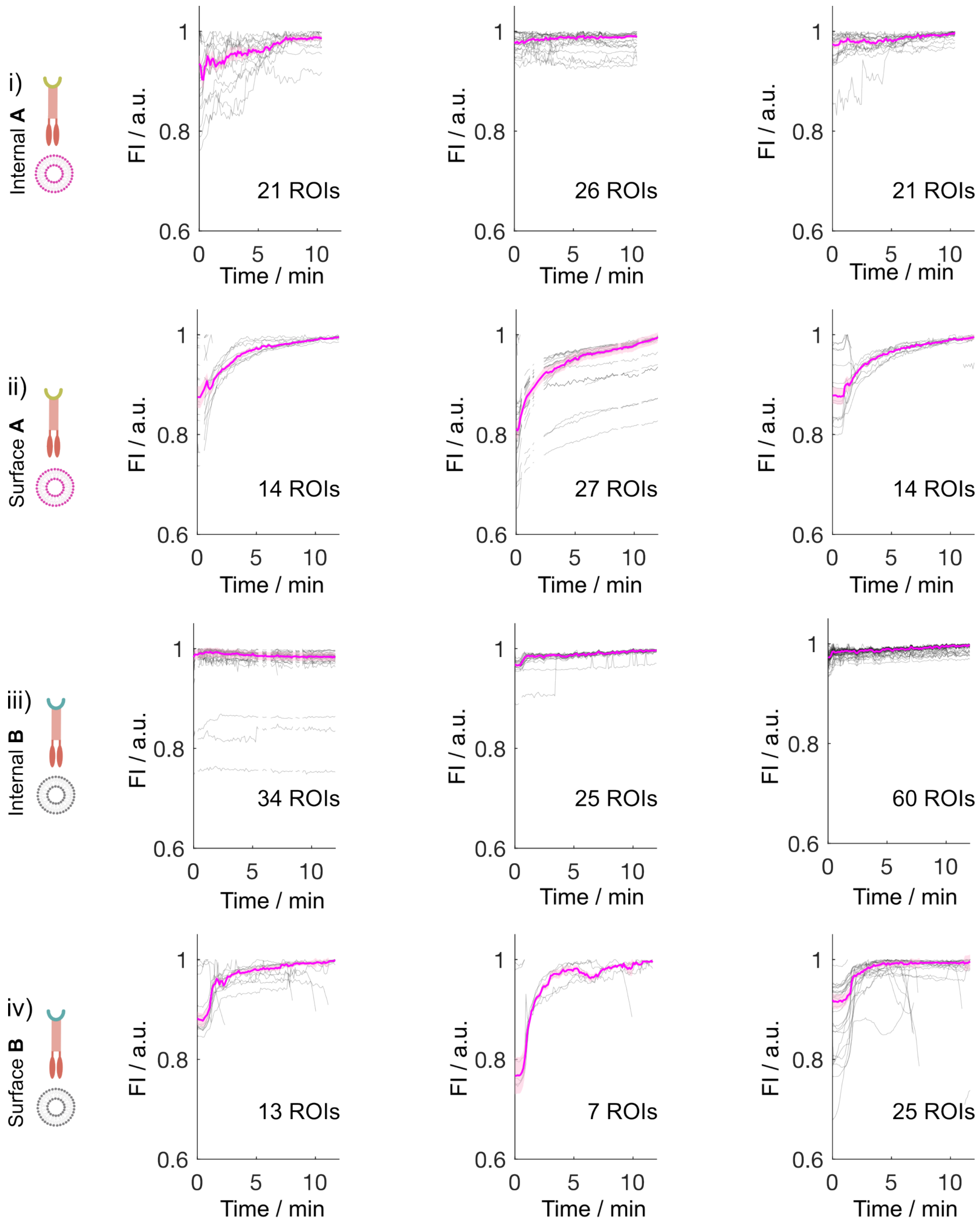


Figure S19: **Fluorescence intensity (FI) of the amphiphilic condensate ROIs normalized by the maximum and monitored over time as LUVs are released and captured by amphiphilic condensates.** The type of LUVs is specified on the left and data are shown for timelapses acquired in three different wells (i.e., three technical repeats). The number of ROIs is based on ROIs identified as shown in Fig. S18. i) Condensates made with NS A, internally-sequestered LUVs, and Cy5-labeled amphiphilic condensate receivers. ii) Condensates made with NS A, surface-tethered LUVs, and Cy5-labeled amphiphilic condensate receivers. iii) Condensates made with NS B, internally-sequestered LUVs, and fluorescein-labeled amphiphilic condensate receivers. iv) Condensates made with NS B, surface-tethered LUVs, and fluorescein-labeled amphiphilic condensate receivers. Release of surface-tethered LUVs shows a larger extent of accumulation of LUVs onto amphiphilic condensates. Discontinuities or sharp drops in signal are caused by frames during which the sample went out of focus.

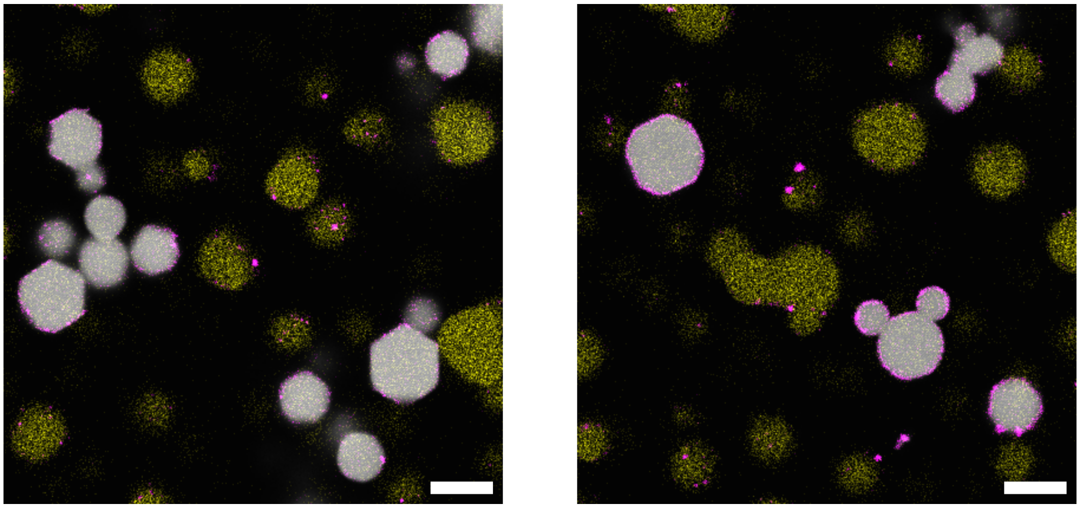

Figure S20: **Two confocal micrographs showing the deposition of LUVs 6 days after release from biphasic DNA sender condensates onto amphiphilic crystalline DNA condensates.** LUVs were initially surface-tethered to B-rich compartments in condensates made with $F_{\mathrm{ab}} = 0.05$. Remaining A-rich compartments are shown in yellow, while amphiphilic receivers are shown in gray (Cy5-labeled). Over time, the remaining A-rich compartments have rounded up. Captured LUVs are shown in magenta. Scale bars: 10 µm.

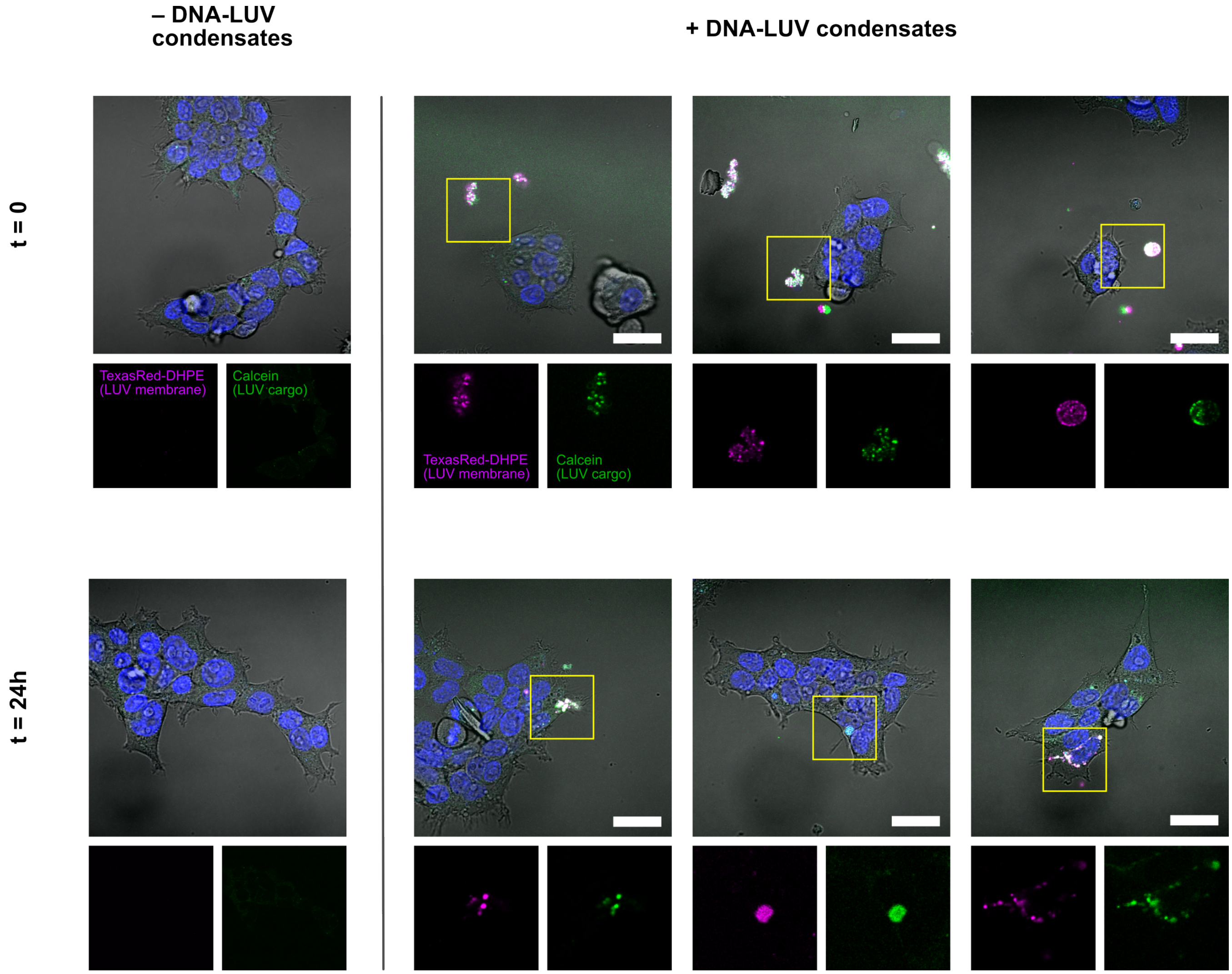


Figure S21: **HEK293 cells can uptake hybrid DNA-LUV condensates over time.** Top row, immediately after addition; bottom row, 24 h later. In each row, the leftmost panel is a negative control showing autofluorescence levels in cells without condensates, and the remaining three are independent fields of view with condensates added. Insets below each fluorescence panel show zoomed Texas Red-DHPE (LUV membrane) and calcein (LUV cargo) regions, contrast-adjusted per inset for easier visualization. Composite overlays show Texas Red-DHPE (magenta), calcein (green), Hoechst (blue), and brightfield (greyscale, 60% opacity). To enable direct visual comparison between samples, display bounds for the calcein, Texas Red-DHPE, and Hoechst channels were set at the 0.5th and 99.9th percentiles of the pooled pixel-intensity distribution across all images and applied identically to every image; the brightfield channel was rescaled to the same percentile bounds computed per image. Condensates were assembled from nanostars A and linkers aa (sequences in Table S1) and annealed without fluorophore-labeled strands owing to limits on the number of dyes that could be imaged. LUVs were labeled with Texas Red-DHPE, loaded with calcein, and incubated with 125 anchors/LUV. Further details are provided in the Methods. Scale bars: 30 µm.

### Supplementary Tables

Table S1: **Oligonucleotide sequences used for the preparation of nanostars and linkers.** Sticky ends (SEs) are emphasized in bold.

| Strand name | Sequence (5' → 3') |
|---|---|
| A4_core1 | **GAT CGC** CGC CGC AAT CAC GCG CGT GCT CGG CGC CAG CAG TCC TGG CG |
| A4_core1_ATTO488 | **GAT CGC** CGC CGC AAT CAC GCG CGT GCT CGG CGC CAG CAG TCC TGG CGT /3ATTO488N/ |
| A4_core2 | **GAT CGC** CGC CAG GAC TGC TGG CGC CGT CGC TTC TCT TCA TAA CAA CG |
| A4_core3 | **GAT CGC** CGT TGT TAT GAA GAG AAG CGT CGC TCT GGC ACA GGT GTA CG |
| A4_core4 | **GAT CGC** CGT ACA CCT GTG CCA GAG CGT GCA CGC GCG TGA TTG CGG CG |
| B4_core1 | **GCG TGT** GCT GTG CAC TGT GAG GAG CGT CGC GTA ACG TTC ATT TGC CG |
| B4_core1_Alexa647 | **GCG TGT** GCT GTG CAC TGT GAG GAG CGT CGC GTA ACG TTC ATT TGC CGT /3AlexF647N/ |
| B4_core2 | **GCG TGT** CGG CAA ATG AAC GTT ACG CGT CGG CGT TGA TCG AGT TAA CG |
| B4_core3 | **GCG TGT** CGT TAA CTC GAT CAA CGC CGT GCA GCG CTC GAC ACA CGT CG |
| B4_core4 | **GCG TGT** CGA CGT GTG TCG AGC GCT GCT CGC TCC TCA CAG TGC ACA GC |
| Linker_aa1 | **GCG ATC** CGC AAA CCA GCA AGC TCA CG |
| Linker_aa2 | **GCG ATC** CGT GAG CTT GCT GGT TTG CG |
| Linker_bb1 | **ACA CGC** CGA CAT GCG TGC GGA GCG CG |
| Linker_bb2 | **ACA CGC** CGC GCT CCG CAC GCA TGT CG |
| Linker_ab_a | **GCG ATC** CGC CTG GTG GAA CGA GCC GC |
| Linker_ab_b | **ACA CGC** GCG GCT CGT TCC ACC AGG CG |
| Linker_aa1_TH | CTG TAC **GCG ATC** CGG CGT CAG ACA GAC GGC CG |
| Linker_aa2_TH | **GCG ATC** CGG CCG TCT GTC TGA CGC CG |
| $\alpha$ trig | CGG CCG TCT GTC TGA CGC CGG ATC GCG TAC AG |
| Linker_bb1_TH | CTC TGA **ACA CGC** CGA CAT GCG TGC GGA GCG CG |
| Linker_bb2_TH | **ACA CGC** CGC GCT CCG CAC GCA TGT CG |
| $\beta$ trig | CGC GCT CCG CAC GCA TGT CGG CGT GTT CAG AG |

Table S2: **Oligonucleotide sequences used for the preparation of cholesterol anchors and docking strands targeting phases A and B, respectively.** The same anchor sequences were used previously by Rubio-Sánchez et al.[9] with inverted directionality. Abbreviations: TEG (triethyleneglycol).

| **Strand name** | **Sequence (5' → 3')** |
|---|---|
| Chol_anchor_1 | **/5CholTEG/** GTT AGT GTG GTG TTT GTG |
| Chol_anchor_2 | CCA AAC AAC AAC ACA ACC CAC AAA CAC CAC ACT AAC **/3CholTEG/** |
| Dock_strand_A | **GAT CGC** GGT TTG TTG TTG TGT TGG |
| Dock_strand_B | **GCG TGT** GGT TTG TTG TTG TGT TGG |
| Dock_strand_linker | **GTC TTC GC** GGT TTG TTG TTG TGT TGG |
| Linker_aa1_anchor | GCG ATC CGC AAA CCA GCA AGC TCA CG |
| Linker_aa2_anchor | GCG AAG ACG CGA TCC GTG AGC TTG CTG GTT TGC G |
| Linker_bb1_anchor | ACA CGC CGA CAT GCG TGC GGA GCG CG |
| Linker_bb2_anchor | **GCG AAG AC** ACA CGC CGC GCT CCG CAC GCA TGT CG |

Table S3: **Statistical data in support of Fig. 3d.** Median values of 2D correlation coefficients ($r_A$ and $r_B$) and median absolute deviation (MAD) calculated over the condensate populations for samples shown in Fig 3**b**. The number of condensates analyzed is given by n $r_A$ and n $r_B$, respectively. As previously discussed, for $F_{ab}$ = 0.05 the analysis was run for the entire network. Stronger LUV localization in the target phases is associated with larger $r_A$ or $r_B$ values.

| **Sample** | $F_{ab}$ | n $r_A$ | $r_A$ (median) | $r_A$ (MAD) | n $r_B$ | $r_B$ (median) | $r_B$ (MAD) |
|---|---|---|---|---|---|---|---|
| **Internal A** | 0.05 | 1 | 0.30 | N/A | 1 | -0.05 | N/A |
| | 0.1 | 134 | 0.38 | 0.02 | 134 | 0.07 | 0.02 |
| | 0.15 | 102 | 0.37 | 0.02 | 102 | 0.14 | 0.06 |
| | 0.2 | 108 | 0.39 | 0.02 | 108 | 0.19 | 0.05 |
| | 0.25 | 179 | 0.34 | 0.02 | 179 | 0.33 | 0.02 |
| | 0.5 | 177 | 0.36 | 0.02 | 177 | 0.37 | 0.02 |
| | 1 | 123 | 0.34 | 0.02 | 123 | 0.34 | 0.02 |
| **Surface A** | 0.05 | 1 | 0.22 | N/A | 1 | -0.03 | N/A |
| | 0.1 | 30 | 0.35 | 0.06 | 30 | 0.19 | 0.03 |
| | 0.15 | 132 | 0.44 | 0.05 | 132 | 0.18 | 0.05 |
| | 0.2 | 69 | 0.54 | 0.04 | 69 | 0.41 | 0.11 |
| | 0.25 | 182 | 0.47 | 0.02 | 182 | 0.45 | 0.03 |
| | 0.5 | 170 | 0.42 | 0.03 | 170 | 0.41 | 0.03 |
| | 1 | 156 | 0.42 | 0.02 | 156 | 0.38 | 0.03 |
| **Internal B** | 0.05 | 1 | -0.05 | N/A | 1 | 0.38 | N/A |
| | 0.1 | 57 | 0.04 | 0.02 | 57 | 0.39 | 0.03 |
| | 0.15 | 141 | 0.09 | 0.04 | 141 | 0.37 | 0.03 |
| | 0.2 | 141 | 0.10 | 0.05 | 141 | 0.35 | 0.03 |
| | 0.25 | 169 | 0.32 | 0.02 | 169 | 0.33 | 0.02 |
| | 0.5 | 211 | 0.31 | 0.02 | 211 | 0.31 | 0.01 |
| | 1 | 151 | 0.34 | 0.01 | 151 | 0.34 | 0.01 |
| **Surface B** | 0.05 | 1 | -0.01 | N/A | 1 | 0.44 | N/A |
| | 0.1 | 71 | 0.18 | 0.06 | 71 | 0.51 | 0.03 |
| | 0.15 | 126 | 0.15 | 0.05 | 126 | 0.50 | 0.04 |
| | 0.2 | 162 | 0.15 | 0.05 | 162 | 0.51 | 0.04 |
| | 0.25 | 118 | 0.53 | 0.03 | 118 | 0.56 | 0.02 |
| | 0.5 | 196 | 0.54 | 0.02 | 196 | 0.57 | 0.02 |
| | 1 | 163 | 0.44 | 0.02 | 163 | 0.43 | 0.02 |

Table S4: **Oligonucleotide sequences used for the preparation of C-star condensates labeled with Cy5 as previously reported by Malouf et al.[10].** Abbreviations: TEG (triethyleneglycol).

| Strand name | Sequence (5' → 3') |
| --- | --- |
| Core1_Cy5 | CGA CGC CGT GAC GCG TTG ATG ACT CGA CTG ACC AG/iCy5/ GCA TCT TAG CTC ACT GGA AAC |
| Core2 | CGA CGC CGT GAC GCG TTT CCA GTG AGC TAA GAT GCT CGA ATG ACT GCA CTG TCA AAC |
| Core3 | CGA CGC CGT GAC GCG TTT GAC AGT GCA GTC ATT CGT CGA ATC GAA ATA CTG TGG AAC |
| Core4 | CGA CGC CGT GAC GCG TTC CAC AGT ATT TCG ATT CGT CTG GTC AGT CGA GTC ATC AAC |
| Terminal chol | GCG TCA CGG CGT CGA A /3CholTEG/ |

Table S5: **Oligonucleotide sequences used for the preparation of C-star condensates labeled with fluorescein as previously reported by Malouf et al.[10].** Abbreviations: TEG (triethyleneglycol).

| Strand name | Sequence (5' → 3') |
| --- | --- |
| Core1_fluorescein | CGA CGC CGT GAC GCG CTT GGG CGT GGC G /iFluorT/ CGC GAG CGC CAA GC |
| Core2 | CGA CGC CGT GAC GCG CTT GGC GCT CGC GTC GGC CAT TGA CTG C |
| Core3 | CGA CGC CGT GAC GCG CAG TCA ATG GCC GTC GGC CAC GCG CAC G |
| Core4 | CGA CGC CGT GAC GCC GTG CGC GTG GCC GTC GCC ACG CCC AAG C |
| Terminal chol | GCG TCA CGG CGT CGA A /3CholTEG/ |
